# Transcription termination factor ρ polymerizes under stress

**DOI:** 10.1101/2023.08.18.553922

**Authors:** Bing Wang, Nelly Said, Tarek Hilal, Mark Finazzo, Markus C. Wahl, Irina Artsimovitch

**Affiliations:** Department of Microbiology and Center for RNA Biology, The Ohio State University, Columbus, OH, USA; Freie Universität Berlin, Institute of Chemistry and Biochemistry, Laboratory of Structural Biochemistry, Takustr. 6, D-14195 Berlin, Germany; Freie Universität Berlin, Institute of Chemistry and Biochemistry, Research Center of Electron Microscopy and Core Facility BioSupraMol, Fabeckstr. 36a, 14195 Berlin, Germany; Helmholtz-Zentrum Berlin für Materialien und Energie, Macromolecular Crystallography, Albert-Einstein-Str. 15, D-12489 Berlin, Germany

## Abstract

Bacterial RNA helicase ρ is a genome sentinel that terminates synthesis of damaged and junk RNAs that are not translated by the ribosome. Co-transcriptional RNA surveillance by ρ is essential for quality control of the transcriptome during optimal growth. However, it is unclear how bacteria protect their RNAs from overzealous ρ during dormancy or stress, conditions common in natural habitats. Here we used cryogenic electron microscopy, biochemical, and genetic approaches to show that residue substitutions, ADP, or ppGpp promote hyper-oligomerization of *Escherichia coli* ρ. Our results demonstrate that nucleotides bound at subunit interfaces control ρ switching from active hexamers to inactive higher-order oligomers and extended filaments. Polymers formed upon exposure to antibiotics or ppGpp disassemble when stress is relieved, thereby directly linking termination activity to cellular physiology. Inactivation of ρ through hyper-oligomerization is a regulatory strategy shared by RNA polymerases, ribosomes, and metabolic enzymes across all life.

## Main

All living cells possess mechanisms that silence expression of potentially harmful or useless DNA. In bacteria, transcription factor Rho/ρ triggers premature release of antisense, horizontally-transferred, and untranslated RNAs^1-3^. ρ is a hexameric, ring-shaped helicase composed of two domains bridged by a flexible connector region^4^. The N-terminal domain (NTD) contains a primary RNA-binding site (PBS), the C-terminal domain (CTD), a secondary RNA-binding site (SBS) and ATPase and helicase modules (Fig. 1a). An open-ring ρ binds to C-rich *rut* RNA sites *via* the PBS and then closes to trap RNA in the SBS, activating ρ ATPase and translocation along the RNA^5-7^. Alternatively, the open ring is recruited to the transcribing RNA polymerase (RNAP) through contacts to RNAP subunits and elongation factors NusA and NusG, then captures the nascent RNA *via* the PBS, and inactivates RNAP^8,9^. Both pathways occur *in vitro*^10^, but ρ trafficking with RNAP in the cell^11^ and the need for ρ targeting to RNAs still in the making, including those that lack *rut* sites, favor the second scenario^12^.

**Fig. 1.**
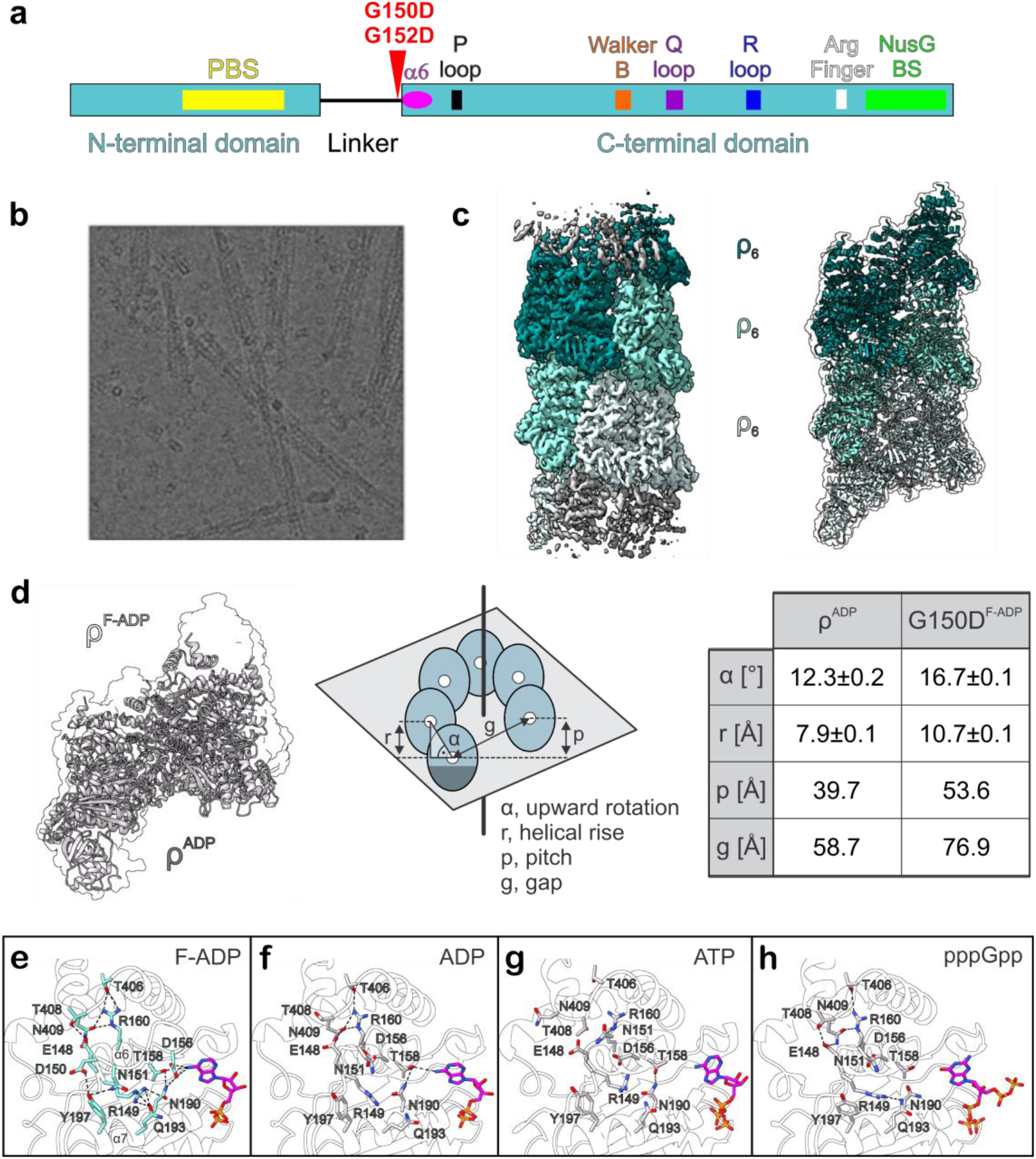
Connector substitutions promote filamentation of *E. coli* ρ. **a**, Schematic diagram of ρ with key regions and positions of filament-inducing substitutions indicated. **b**, Negatively stained *in vitro* ρ G150D filaments. **c**, CryoEM reconstructions of G150D filaments. **d**, Comparison of WT and G150D hexamer geometries. Values of protomer-protomer distances and angles are derived from the draw_rotation axis script in Pymol. **e-h**, G150D- and nucleotide-induced changes in the interaction network of α5/α6 loop (residues 149-156) and helix α6 (residues 155-166). F-ADP, ADP-bound ρ G150D filament. See Extended Data Figs. 1 and 2 for cryoEM analysis.

ρ surveils the nascent RNA to ensure that only those RNAs that are either translated by the ribosome or protected by dedicated anti-termination factors^1^ are made. However, during slow growth or stress, when translation is inefficient, indiscriminate termination by ρ may be lethal; consistently, partial loss-of-function mutations in the *rho* gene enable *E. coli* survival on ethanol, which inhibits translation^13,14^. ρ cellular levels are subject to autoregulation^15^, suggesting that ρ must be inhibited during stress, *e*.*g*., by accessory proteins that inhibit ρ-RNA interactions^16,17^.

## Results

### ρ connector mutants form filaments

Studies of poorly translated xenogeneic operons may reveal mechanisms of cellular adaptation to translational stress. The cell wall biosynthesis *waa* operon is silenced by ρ unless an antitermination factor RfaH is present^18^. Using genetic selection for Δ*rfaH* suppressors, we identified two unexpected changes in the ρ connector encoding G150D and G152D variants defective in termination (Fig. 1a). We hypothesized that G-to-D substitutions may rigidify the connector, disrupting communications between ρ domains and inhibiting ring closure^18^. We thus determined structures of G150D and G152D ρ proteins using cryogenic electron microscopy (cryoEM). Strikingly, we found that both variants formed extended helical filaments (Fig. 1b and Extended Data Fig. 1) that trap ρ in an inactive state unable to interact with either RNA or RNAP; in the main text, we focus on G150D.

Analysis of cryoEM structures of G150D filaments reveals that 18 ρ protomers are well-defined in the density even though 2D micrographs show that the filament extends far beyond (Fig. 1b, c). We prepared G150D in the presence of the ATP transition-state analog ADP•BeF_3_ but, while the density for ADP is visible in the ATP-binding pocket at each interface, the BeF_3_ moiety is absent. The filament forms along a left-handed helical axis, similar to the wild-type (WT) ρ-hexamer in an open-ring conformation reported by Skordalakes and Berger^4^, who proposed that an increase in pitch of the ρ ring would foster oligomerization. Consistently, in the G150D structure, the helical pitch increases, showing an 11 Å rise between subunits and a pitch of 17°, compared to 8 Å and 12° in WT ρ (Fig. 1d). This leads to an enlarged opening of the ring, thus allowing an additional hexamer to join the ring.

By progressing into a filamentous structure, ρ G150D forms additional contacts between its NTD and CTD. The filament is strengthened by interactions of the very C-terminal α-helix (α16; residues 408-418) of one subunit (ρ_1_) with the NTDs of protomers ρ_7_ and ρ_8_ located seven and eight subunits upstream the helical axis, respectively. The last three α helices (α14-α16) of the ρ_1_ CTD are inserted between the central (loop β1/β2-β2-β3-α4, residues 59-88) and C-terminal (loop β4/β5, residues 103-110) portions of the ρ_7_ and ρ_8_ NTDs, respectively. R87 of ρ_7_ reaches towards the loop between α15 and α16 of ρ_1_ and makes contacts to M405 and M415, while the backbone carbonyl of D60 in ρ_7_ forms a hydrogen bond to K402. Likewise, E106 of ρ_8_ approaches K379 and K417 within α14 and α16 of ρ_1_, respectively.

Residue G150 is located within α5/α6 loop (residues 149-156) sandwiched between helix α7 (residues 184-198 that connect to the P-loop of the ATP pocket) and the helix-loop-helix region formed between α15 and α16. The α5/α6 loop is followed by helix α6 (residues 155-166), which not only faces the nucleobase of the bound nucleotide but also interacts with residues in α14-α16 (Fig. 1e). Structural comparison to WT ρ bound to ADP (Fig. 1f) revealed that the G150D substitution induces major rearrangements of the intra-molecular network within α5/α6 loop and proximal regions without affecting the overall structure of the protomer. In G150D, a partial unwinding of α6 at its N-terminal end repositions several residues to form new interactions. D156, which points away from the nucleotide in WT ρ, now directly forms a hydrogen bond with the nucleobase of the bound ADP and contacts N190 within α7. In addition, D150 forms a hydrogen bond to Y197. Thus, the G150D/G152D substitutions seem to restrict an otherwise very flexible arrangement of the nucleotide-binding site (Fig. 1g, h) by tethering α5/α6 loop more tightly to neighboring regions of the CTD and to the bound nucleotide.

### Wild-type ρ forms filaments *in vitro* and *in vivo*

To test whether purified WT and G150D ρ proteins form filaments in solution, we first used pelleting assays, in which protein filaments sediment at 100,000 *g*, whereas monomers and smaller oligomers remain in the supernatant. As observed for ParM filaments^19^, we found most G150D ρ in the pellet, whereas WT ρ remained soluble (Fig. 2a). Nucleotide cofactors are known to affect the formation of protein filaments^19,20^, and the loss of BeF_3_ suggests that G150D filaments preferentially bind ADP (Fig. 1e). We found that ADP promoted, while ATP and ATP-γS slightly inhibited, aggregation of the WT ρ, but had little effect on the G150D variant (Fig. 2a).

**Fig. 2.**
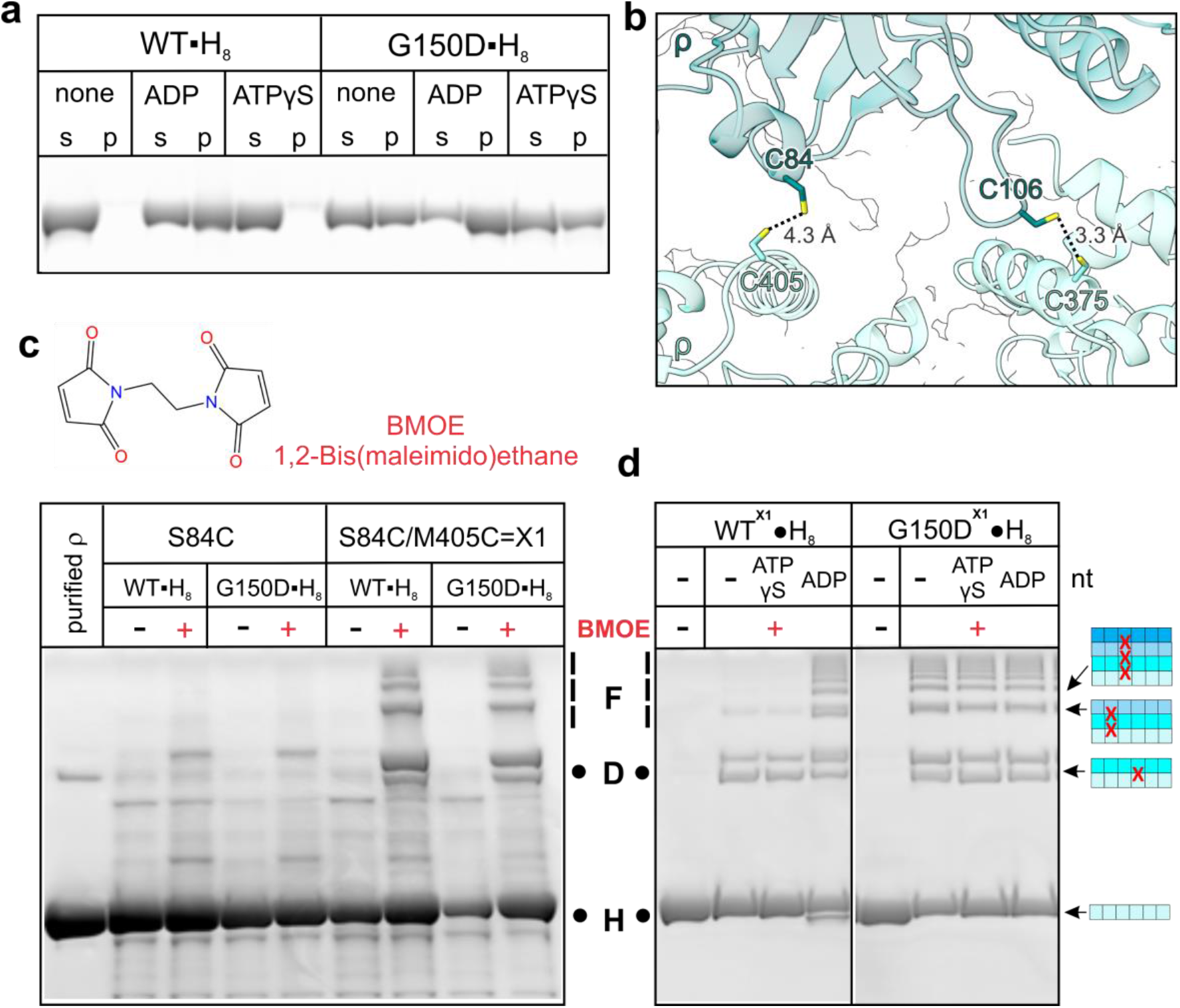
Filament detection by pelleting and cross-linking. **a**, Nucleotide-dependent polymerization of ρ. 2 μM WT or G150D ρ were incubated with nucleotides (2 mM) and centrifuged at 100,000 *g* for 20 min at 20 °C. Supernatant (s) and solubilized pellets (p) were analyzed by LDS–PAGE and Coomassie Blue staining. **b**, Sensor Cys pairs, C84/C405 and C106/C375, at the filament interface. **c**, BMOE-mediated *ex vivo* cross-linking of *E. coli* strains expressing WT or G150D ρ with C84/C405 residues. Reaction products were detected by LDS-PAGE and in-gel fluorescence using His-tag specific NTA-ATTO 550 stain. **d**, BMOE-mediated cross-linking of purified 1 μM WT^X1^•H_8_ and G150D^X1^•H_8_ ρ in the presence of 2 mM nucleotides (nt) or water. Reactions were analyzed as in (**c**). Cartoons depict linearized single and stacked ρ rings that, when crosslinked (a red X) *via* the engineered Cys residues at the interface, can give rise to the observed products. Each ρ subunit can only be crosslinked to two neighbors, one from the ring above, one from the ring below. Thus, a species that migrates as a dimer corresponds to at least two stacked ρ rings. H, D, and F indicate hexamers, dodecamers, and filaments, respectively.

Pelleting assays cannot distinguish between filaments and large amorphous aggregates. To test whether ρ forms filaments as revealed by the structures, we designed “sensor” Cys residues that would be closely spaced only in filaments (Fig. 2b) and used bismaleimidoethane (BMOE), a high-efficiency short-length (8.0 Å) cell-permeable sulfhydryl-to-sulfhydryl crosslinker, to assess filament formation^21^. We found that crosslinks readily formed *ex vivo* when BMOE was added to intact cells expressing C-terminally octahistidine-tagged (H_8_) WT or G150D ρ with C84/C405 (X1; Fig. 2c) or C106/C375 (X2; Extended Data Fig. 3) substitutions from plasmids. The crosslinks were observed only in the presence of BMOE and both Cys residues and formed a pattern consistent with a mixture of dodecamers and higher-order oligomers. We did not observe differences between the WT and G150D variants, possibly due to very high levels of expression used to outcompete abundant chromosomally-encoded ρ.

**Fig. 3.**
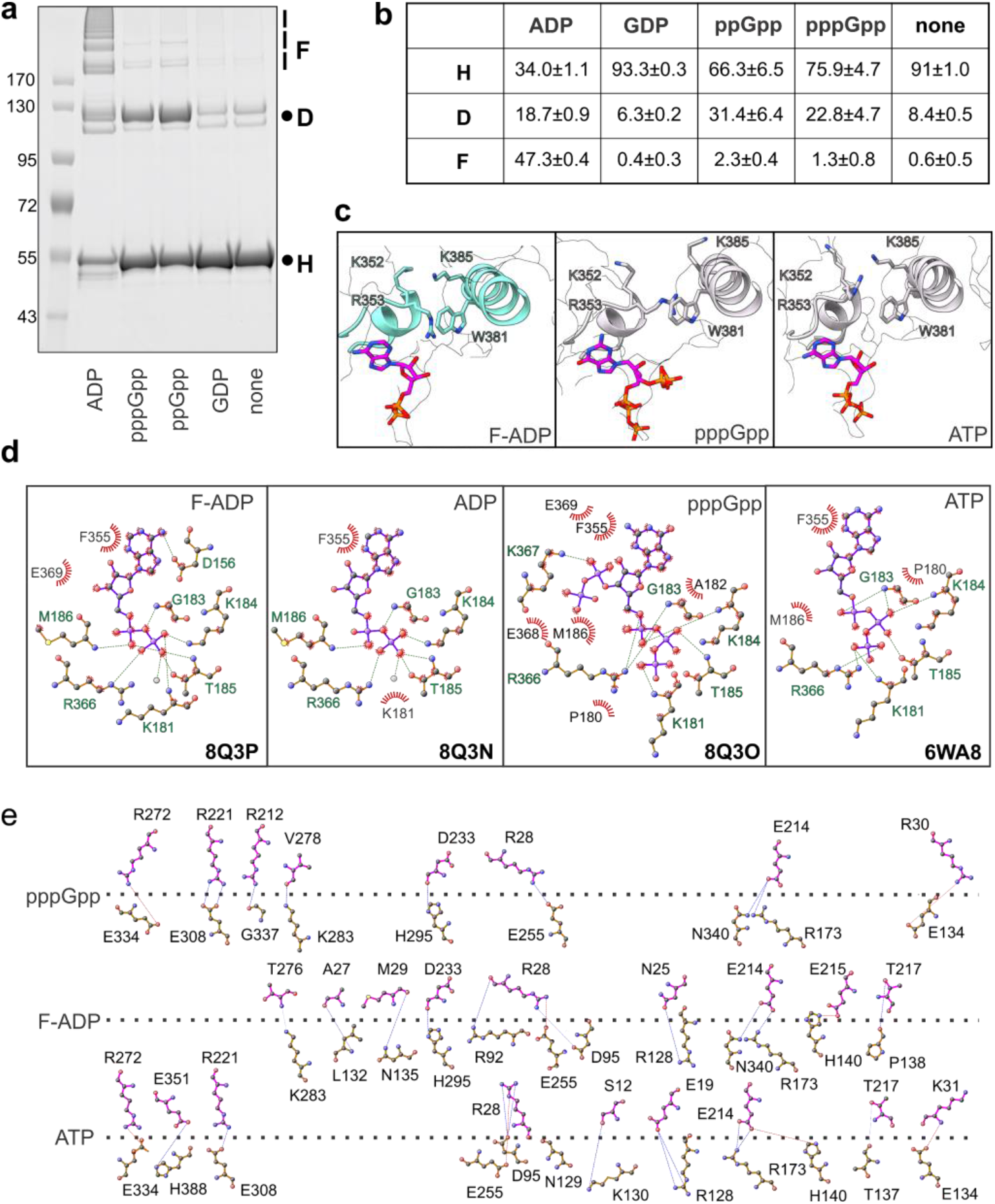
ADP and (p)ppGpp promote hyper-oligomerization by changing ρ protomer interfaces. **a**, Detection of ρ oligomers by BMOE crosslinking in the presence of ADP, GDP, and (p)ppGpp; protein marker sizes (in kDa) are indicated on the left. **b**, Distribution of oligomeric states induced by different nucleotides reported as mean ± SD, n=3. **c**, ρ G150D-ADP and ρ WT-pppGpp structures show a strong cation-π interaction between R353 and W381, whereas in the WT-ATP structure the two residues are more flexible. **d**, LigPlot representation of the nucleotide-binding modes of ρ. Residues involved in binding to the indicated nucleotides are labelled. Green dashes, hydrogen bonds; red rays, van der Waals interactions. **e**, DimPlot representation of interactions across the ρ B/C protomer interface shown by a horizontal dashed line. Green dashes, hydrogen bonds; red dashes, salt bridges.

Crosslinking of purified ρ variants *in vitro* showed that G150D^X1^•His_8_ formed oligomers in the absence of nucleotides, whereas WT ^X1^•His_8_ was crosslinked only in the presence of ADP (Fig. 2d), mimicking the results of pelleting assays (Fig. 2a). In the gel, we observed ρ monomers (∼ 55 kDa), dimers, and higher-order oligomers. To simplify the description, we assign ρ monomers to hexamers (H) and dimers to dodecamers (D), although they can represent smaller assemblies, *e*.*g*., pentamers and 11-mers, respectively; we term all larger species filaments (F). A ladder of crosslinked monomers, up to at least nine species that correspond to 8+ complete ρ rings, is visible on the gel, whereas larger species (500+ kDa) could not be resolved. In each case, most easily seen with the dimers, we observed two species with different mobilities, which we presume arise due to differences in LDS binding.

In these experiments, we used His-tagged proteins to enable visualization in cell extracts using a sensitive NTA-ATTO stain. While the ρ C-terminus is not conserved (Extended Data Fig. 4a), and the tag does not alter ρ activity *in vitro* (data not shown), two observations made us wonder if the tag could induce filament formation: (i) we did not detect filaments on grids with tag-less proteins and (ii) we observed a density attributable to the tag near the filament interface (Extended Data Fig. 4b). Our findings that the tag-less WT ρ behaved similarly to its His-tagged counterpart in pelleting assays and that WT ^X1^ and G150D^X1^ were crosslinked irrespective of the tag (Extended Data Fig. 4c,d) argue that the tag is not required for filament formation when ρ is present at 1 μM, its cellular concentration^22^. Nonetheless, G150D^X1^ • His_8_, but not G150D^X1^, formed filaments at 50 nM (Extended Data Fig. 4e), explaining why we observed filaments only with tagged G150D and G152D proteins on grids, when low concentrations are used. Subsequent experiments were carried out with tag-less proteins, unless noted otherwise.

**Fig. 4.**
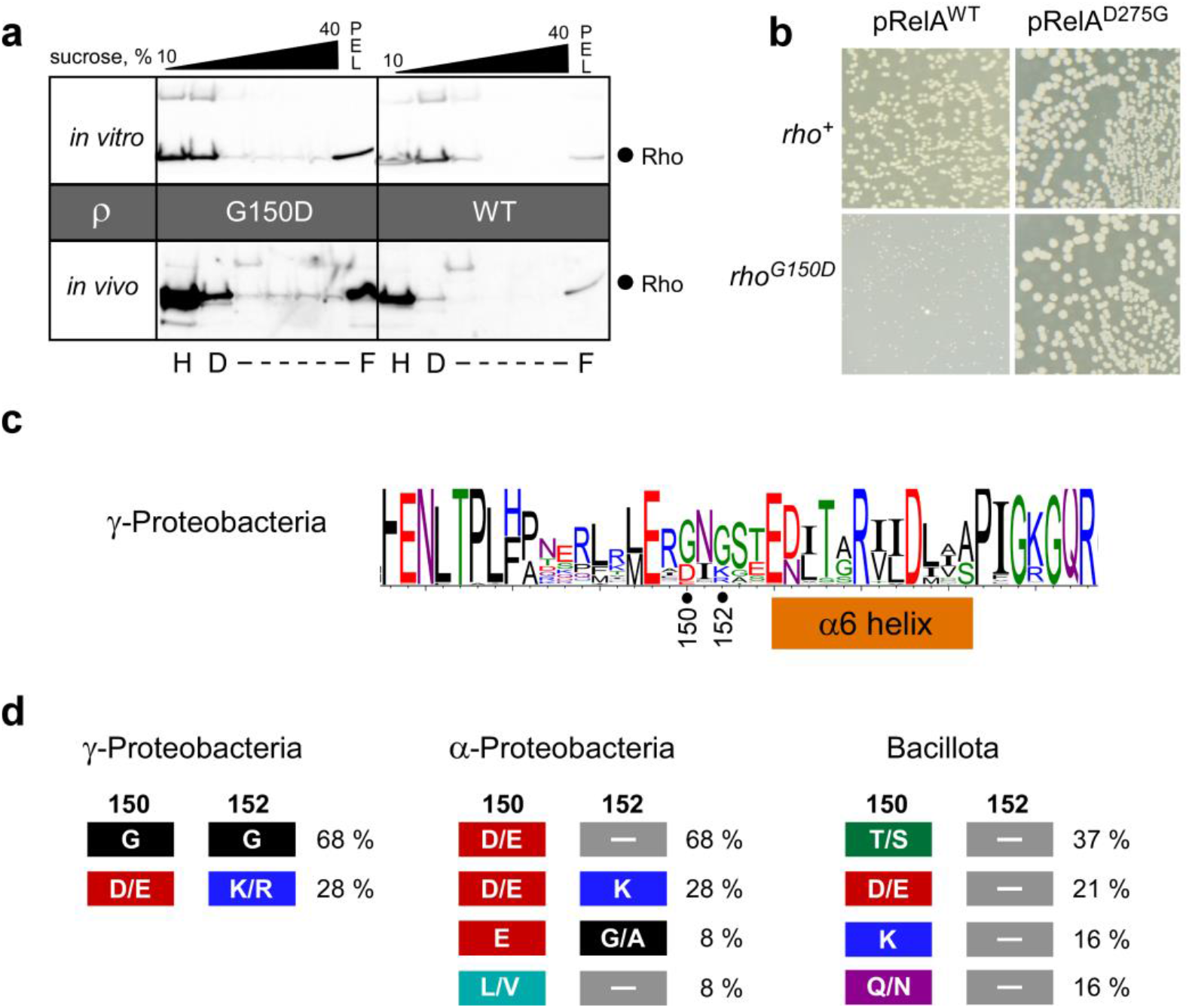
ρ G150D forms oligomers and is hypersensitive to ppGpp in the cell. **a**, Sucrose density gradient analysis of WT and G150D ρ proteins *in vitro* (top) and in cells (bottom). ρ was detected by Western blotting with polyclonal anti-ρ antibodies. H, D, and F positions were assigned based on sedimentation of known proteins. **b**, WT and G150D *rho* cells were transformed with plasmids expressing either WT or D275G *relA* and plated on LB with carbenicillin in the absence of induction. **c**, Residue conservation around the connector and α6 helix in γ-Proteobacteria. Positions 150 and 152 are indicated. **d**, Variation of residues corresponding to G150 and G152 in *E. coli* ρ. The percentages of different combinations are calculated for each cluster: γ-Proteobacteria (cluster 1), α-Proteobacteria (cluster 2), and Bacillota (cluster 7). The relative positions for 150 and 152 were identified from a multiple sequence alignment generated from our representative sequences; see Methods for more details and dataset 1 for the NCBI sequence IDs.

### (p)ppGpp binds to ATP site and induces ρ oligomerization

While ADP promotes ρ filament formation *in vitro* (Fig. 2a, d), ADP levels do not change dramatically during stress^23^, questioning the physiological relevance of this observation. We wondered whether ρ could directly sense the stress alarmone (p)ppGpp^24^ that can be crosslinked to ρ^25^. ppGpp quenches fluorescence of a unique W381 residue, strongly suggesting that ppGpp binds to the same site as ATP at the interface between ρ subunits (Extended Data Fig. 5a, b). We found that ppGpp and pppGpp, but not GDP, promoted ρ hyper-oligomerization (Fig. 3a), but favored the formation of smaller oligomers, as compared to ADP. With ppGpp, almost a third of crosslinked ρ was present in the dodecamer fraction, and only ∼2 % formed short filaments. By contrast, with ADP, almost half of ρ was in the filament form (47 %) and dodecamers accounted for ∼19 % (Fig. 3b).

**Fig. 5.**
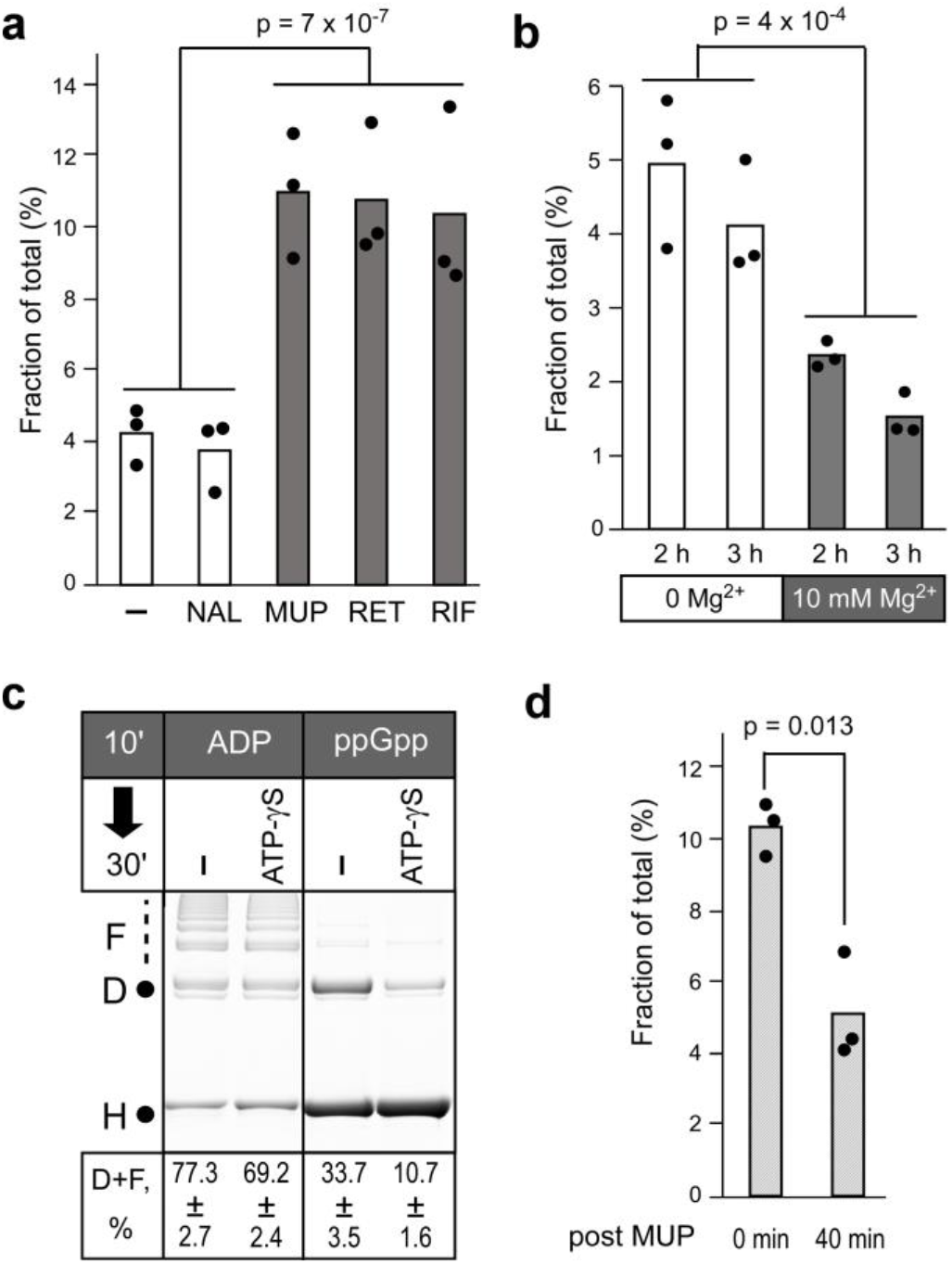
ρ oligomers form and disperse in response to cellular cues. **a**, ρ forms large oligomers following a 30-min exposure to antibiotics that inhibit protein or RNA synthesis. ─, none; NAL, nalidixic acid; MUP, mupirocin; RET, retapamulin; RIF, Rifampicin. The p-value was calculated between (─, NAL) and (MUP, RET, RIF). **b**, Cellular starvation for Mg^2+^ leads to formation of ρ polymers. Cells were grown for 2 h and 3 h in modified MOPS medium supplied with either 0 mM or 10 mM MgCl_2_. **c**, 2 mM ADP- and ppGpp-stabilized ρ polymers were performed *in vitro* for 10 min and challenged with a non-hydrolysable ATP analog (20 mM) for 30 min, followed by BMOE crosslinking. Percentage of ρ found in the D (dodecamer) and F (filament) fractions is shown as mean ± SD; n = 4. **d**, Following MUP treatment (as in panel **a**) cells were washed and incubated for 40 min in the absence of MUP (before growth resumes; Extended Data Fig. 7. Two-tailed T-test assuming unequal variance was applied for p-value calculation. Fraction of total (%) represents the amount of ρ pelleted at 100,000 *g*. See Methods and Extended Data Fig. 7 for details of experiments in panels **a, b**, and **d**.

An apparent distribution of ρ oligomeric states reflects their presence in solution and efficiency of crosslinking by BMOE and may be altered by Cys residues. To visualize oligomers of WT “native” ρ, we used sucrose gradient centrifugation. We observed that, in the absence of nucleotides, WT ρ was present in two fractions, consistent with a mixture of hexamers and dodecamers (Extended Data Fig. 5c), as reported previously^26,27^. In the presence of ppGpp and ADP, ρ distribution was shifted toward progressively longer oligomers, in agreement with our crosslinking assays.

The differences between ADP and ppGpp may be due to differences in binding affinities or to structural changes induced upon the nucleotide binding to ρ. Crosslinking suggested that ppGpp binds to ρ weakly *in vivo*^25^; consistently, we were unable to accurately measure ppGpp affinity using ITC, DRACALA or fluorescence quenching assays. In fact, the low binding affinity could safeguard ρ from fortuitous inactivation, except during acute stress, when ppGpp can reach mM concentrations^23^. To investigate the effects of (p)ppGpp on ρ, we solved the cryoEM structure of ρ bound to pppGpp; ppGpp did not stably bind to ρ under conditions of grid preparation. Structural analysis using single-particle cryoEM revealed one reconstruction of an open ρ hexamer with pppGpp bound at four ATP-binding pockets. Overall, the structure resembles other open-ring ρ structures except for residues that form the ATP pocket, with changes to the intra-molecular network within the loop region and helices α6 and α7 (Fig. 1g, h). Here, T158 forms a hydrogen bond to N190 while E155 is engaged in an electrostatic interaction with R149. In contrast, in the ADP-bound state, T158 interacts with the amino group of the nucleobase and R149 hydrogen bonds to both N190 and Q193.

The ρ ring dynamics is thought to be controlled by a molecular switch involving residues at the protomer interface^28^. An inhibitory cation-π interaction between R353 and W381 has been suggested to stabilize the open conformation and prevent a productive contact between R366 (the Arg finger) and ATP^28^. The R353-W381 cation-π interaction is similar in the pppGpp- and F(filament)-ADP-bound states and identical between neighboring protomers (Fig. 3c). In contrast, in the ATP-bound state, the R353-W381 contact is more dynamic, as we observe distinct orientations of these two residues within the hexamer. The cation-π interaction must be disrupted during ring closure^28^, and higher flexibility of R353 and W381 likely facilitates the underlying conformational changes.

We carried out a detailed comparison of the binding modes of nucleotides^29^ to WT hexamers and G150D filaments (Fig. 3d). In the pppGpp-bound structure, we observed additional contacts of the 3’-pyrophosphate, with the δ-phosphate forming a hydrogen bond to the amino group of K367 and van der Waals interactions to E368 and E369. The nucleotide is further stabilized by G183 and K184 of one protomer and by R366 at the interface forming a hydrogen bond with the α-phosphate. Additionally, T185 forms a hydrogen bond with the β-phosphate, while K181 contacts the γ-phosphate. In contrast, in the ATP-bound structure, the α-phosphate only forms a single hydrogen bond with the peptide backbone of G183, and R366 exclusively interacts with the γ-phosphate. Like pppGpp, ADP is bound to G150D and WT ρ proteins *via* an extensive interaction network involving the α- and β-phosphates.

We also observed differences in the interaction networks at the interfaces between protomers depending on the nucleotide bound (Fig. 3e). Compared to WT/ATP, G150D/ADP exhibits more interactions along the entire protomer interface, involving the NTD, the connector, and the CTD. Both G150D/ADP and WT/pppGpp exhibit slightly more contacts between their CTDs compared to WT/ATP, while in WT/pppGpp there are significantly fewer interactions along the NTDs and the connector regions. The resulting nucleotide-specific alignment of the protomers likely accounts for the geometry of the ρ hexamer and thus the potential to form higher oligomeric states. Altogether, our structural analysis suggests that when ρ is bound to ADP or pppGpp, the interaction network within the nucleotide-binding pocket is strengthened, in turn leading to changes at the protomer-protomer interfaces and enabling ρ to form an open hexamer with a higher helical pitch (Fig. 1d).

### G150D substitution promotes hyper-oligomerization and sensitivity to ppGpp *in vivo*

Unlike WT ρ, the G150D variant forms filaments in the absence of ADP or (p)ppGpp *in vitro* (Fig. 2a). To compare cellular oligomeric states of WT and G150D, we fractionated total cell extracts from strains carrying the WT or G150D *rho* chromosomal alleles on sucrose gradients; purified proteins were used as controls (Fig. 4a). We found that WT ρ was present predominantly as a hexamer in the cell, whereas G150D formed higher-order oligomers that were distributed along the gradient, with a substantial fraction (presumably long filaments) present in the pellet (Fig. 4a). These results confirm that G150D ρ Polymerizes even in the absence of stress and suggest that it could be hypersensitive to (p)ppGpp. Consistently, we found that the growth of the G150D strain was inhibited by a plasmid-borne ppGpp synthetase RelA, but not its catalytically inactive D275G variant, whereas the WT strain was resistant to both (Fig. 4b). Conversely, the deletion of *relA* promoted growth of G150D, but not of the WT strain (Extended data Fig. 5d). Finally, increased production of (p)ppGpp triggered by mupirocin (MUP)^23^, an inhibitor of isoleucyl-tRNA synthetase, strongly inhibited G150D, but had lesser effects on the WT strain and a mutant with reduced levels of ρ (Extended Data Fig. 5e).

G150D partially compromises ρ activity^18^ but some proteobacterial proteins have Asp at this position (Fig. 4c). To solve this puzzle, we revisited the phylogenetic analysis of ρ, last carried out a decade ago in a smaller sequence space^30^. Our analysis confirmed that residues implicated in mechanochemistry and RNA binding are conserved, whereas the connector and regulatory elements are divergent (Extended Data Fig. 6). Residues at positions 150 and 152 are conserved only in γ-Proteobacteria and fall into two classes. While most ρ proteins have Gly at both positions, 28 % have Asp/Glu at 150 and Arg/Lys at 152 (Fig. 4d), a covariation that is likely to avoid the disruptive effects of the sole negative charge observed here.

**Fig. 6.**
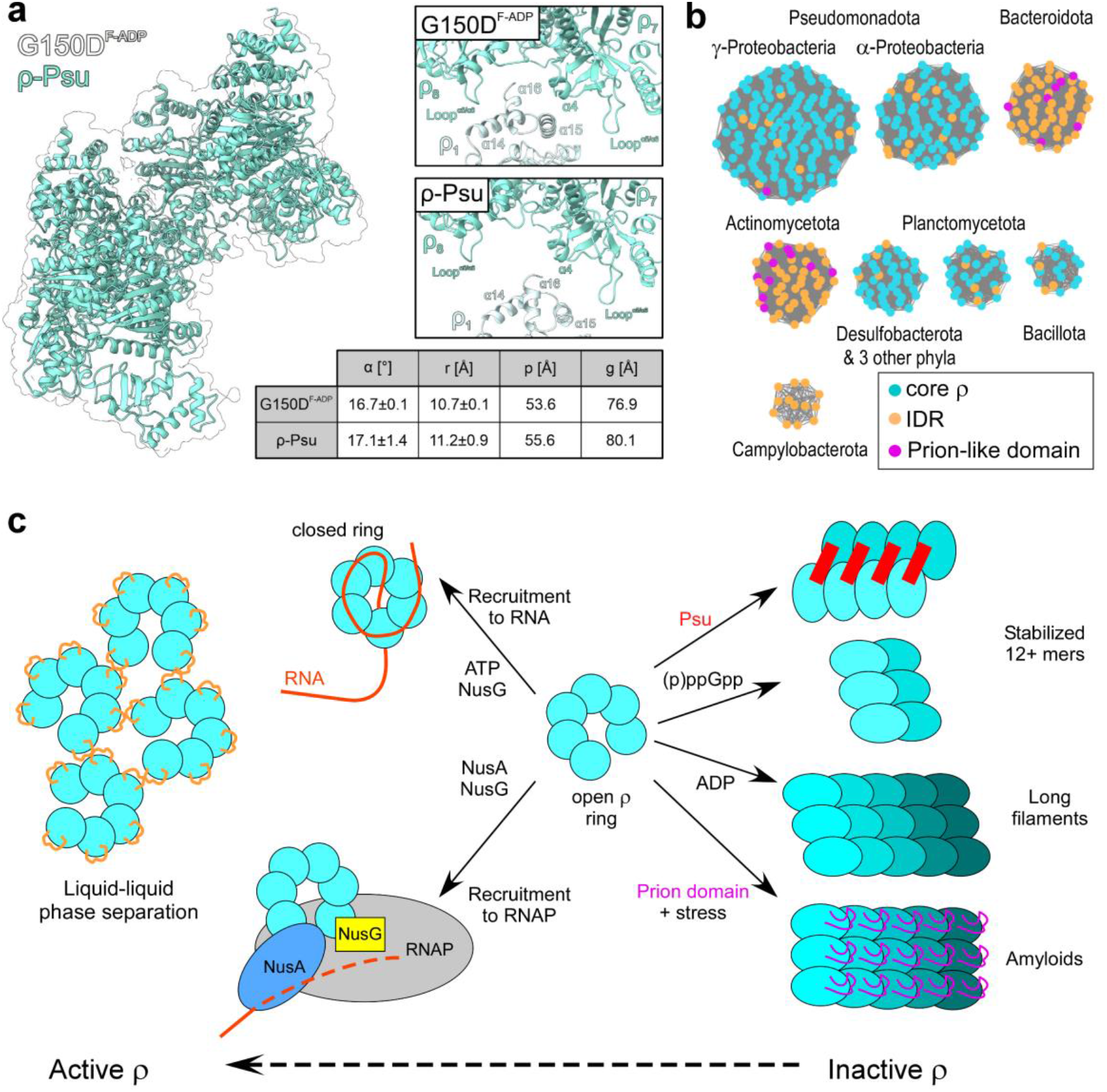
Controlling ρ oligomeric state. **a**, Comparison of ρ hexamers extracted from ADP-stabilized G150D filaments (surface) and from expanded ρ complexes locked by Psu (cartoon; for details, see accompanying manuscript). Detailed views show a comparison of the stacking of ρ subunits in ADP-stabilized G150D filaments and expanded ρ complexes locked by Psu. Helical parameters as defined in Fig. 1d. **b**, Eight major clusters of ρ proteins determined by Markov Clustering algorithm based on sequence similarity; nodes represent ρ proteins, and pair of nodes are connected by edge (grey line) if the similarity score is highly confident (e-value < 10^−99^); hypothetical IDRs (intrinsically disordered regions) and prion domains are indicated. **c**, Model for different modes of ρ aggregation (ordered polymerization/filamentation, phase separation or amyloid formation); while polymerization/filamentation and amyloid formation inactivate ρ, phase separation can activate ρ.

### Stress-induced ρ hyper-oligomerization is reversible

We conjectured that ρ may form inactive oligomers during translational stress, when RNA becomes unprotected, or when RNA synthesis is arrested, triggering ρ release from RNAP. To test this hypothesis, we analyzed the ρ oligomeric state upon exposure of exponentially-growing *E. coli* cells to antibiotics (Fig. 5a). To stop protein synthesis, we used retapamulin (RET), which arrests the initiating ribosome^31^ and MUP, which induces ribosome stalling during elongation^32^. To inhibit RNA synthesis, we used rifampicin (RIF), which blocks promoter escape^33^. As a control, we used nalidixic acid (NAL), an inhibitor of DNA gyrase^34^. We found that ρ was enriched in the pellet (filament) fraction following 30-min exposure to MUP, RET, and RIF, but not after NAL or mock treatment (Fig. 5a). We also observed that ρ partitions into the pellet under Mg starvation (Fig. 5b), which promotes (p)ppGpp accumulation^35^.

Are ρ oligomers dead-end complexes or transient dormant states that dissociate into active hexamers when stress is relieved? To test whether filaments could be dispersed *in vitro*, we pre-incubated WT ρ with ADP or ppGpp, followed by the addition of ATP-γS for 30 min (Fig. 5c). We observed that ADP-stabilized filaments were only partially perturbed, whereas two-thirds of ppGpp-stabilized oligomers disappeared when challenged with ATP-γS. These results suggest that ppGpp-ρ oligomers are in equilibrium with ρ hexamers, while ADP-bound filaments are more stable, consistent with our analysis of the protomer interfaces (Fig. 3e). We also found that ρ oligomers triggered by MUP, which induces synthesis of (p)ppGpp^23^, can be dispersed upon antibiotic removal (Fig. 5d). Collectively, our results show that *E. coli* ρ can reversibly hyper-oligomerize in response to cellular cues.

## Discussion

Our results show that under stress, termination factor ρ can be temporarily inactivated by sequestration in higher-order polymers and filaments. Intriguingly, phage P4 apparently capitalizes on the intrinsic flexibility of the open ρ ring to implement a similar strategy of ρ inactivation during host takeover. Multiple copies of the dimeric P4 capsid protein, Psu, can directly bind two open ρ rings, stabilizing a ρ helical geometry very similar to the conformation observed in the nucleotide-induced ρ polymers reported here and, thus, facilitating expansion of the rings to at least the nonamer stage (see accompanying manuscript; ^36^; Fig. 6a). However, stacked ρ subunits in the Psu-stabilized, expanded rings are not as snuggly interdigitated as in the nucleotide-induced ρ polymers (Fig. 6a, right). Irrespectively, ρ hyper-oligomerization is obviously a versatile strategy to inactivate ρ in diverse stress situations.

Biomolecular phase separation covers a continuum of states, from dynamic liquid droplets held together by weak and transient interactions to highly ordered stable polymers^37,38^. Bacterial ρ proteins apparently utilize this entire spectrum to control their activity (Fig. 6). While all ρ proteins share features that promote phase separation, *e*.*g*., multivalence and RNA binding^39^, our clustering analysis shows that intrinsically-disordered regions (IDRs), which mediate weak interactions in liquid droplets^40^, are rare in Bacillota and Pseudomonadota but common in Bacteriodota and Actinomycetota (Fig. 6b), and the residues at the protomer interfaces are variable (Extended Data Fig. 8. We propose that phylum-specific features of ρ proteins underpin their different survival strategies (Fig. 6c). A recent study showed that an IDR in *Bacteroides thetaiotaomicron* ρ mediates formation of liquid condensates that promote ρ activity and survival in the gut^41^. By contrast, in *C. botulinum*, a prion-like IDR fosters the formation of rigid amyloid aggregates in which ρ is inactive, a transition thought to promote adaptation to stress^42^. Here, we show that *E. coli* (Fig. 2a, 3a) and *Pseudomonas aeruginosa* (Extended Data Fig. 9) ρ proteins, which lack prion-like domains or IDRs, form higher-order oligomers. We conjecture that the formation of trapped, yet reversible, polymers is a cost-efficient mechanism by which cells temper ρ activity under acute stress. Unlike aggregates, which are likely disassembled by proteolysis, ρppGpp oligomers can disassemble under conditions that signal the return to optimal growth (Fig. 5c, d). The reversible sequestration obviates a need for *de novo* protein synthesis when the stress is relieved, and stress-induced hibernation by multimerization is a common adaptive response shared by ribosomes, RNAPs, and metabolic enzymes in bacteria and eukaryotes^43-47^. In contrast, hyper-stabilization by Psu protein^36^ is lethal in many pathogens^48^, suggesting that ADP- and Psu-like ligands may be attractive drug leads.

## Data availability

CryoEM reconstructions have been deposited in the Electron Microscopy Data Bank (https://www.ebi.ac.uk/pdbe/emdb) and structure coordinates have been deposited in the RCSB Protein Data Bank (https://www.rcsb.org). Accession codes are listed in Table S1.

## Acknowledgements

We thank Georgy Belogurov, Jan Löwe, Natacha Ruiz, and P. Lydia Freddolino for discussions, Evgeny Nudler for anti-ρ antibodies, and Mike Laub for RelA-expression plasmids. This work was supported by grants from the National Institutes of Health (GM067153 to I.A.), the Deutsche Forschungsgemeinschaft (INST 130/1064-1 FUGG to Freie Universität Berlin; WA 1126/11-1, project number 433623608, to M.C.W.), and the Berlin University Alliance (501_BIS-CryoFac to M.C.W.). We acknowledge the assistance of the core facility BioSupraMol supported by the Deutsche Forschungsgemeinschaft in electron microscopic analyses and the Sanger Sequencing at the OSU Comprehensive Cancer Center supported in part by NCI P30 CA0168058.

## Author contributions

B.W. analyzed filament formation using *in vitro* and *in vivo* approaches and performed bioinformatic analyses. N.S. assembled complexes for structural analysis and built atomic models. T.H. acquired, processed and refined cryoEM data. M.F. performed growth assays. I.A. constructed plasmids for protein expression. I.A. wrote the first draft with contributions from N.S. and B.W. All authors contributed to data interpretation and manuscript revisions.

