## Supplementary material for "Transcription termination factor ρ polymerizes under stress"

### Experimental Procedures

#### Protein purification

Plasmids used for protein purification are listed in Table S2. All of the restriction and modification enzymes for plasmid construction were from New England Biolabs. DNA oligonucleotides for vector construction and sequencing were obtained from Millipore Sigma, and synthetic DNA fragments for Gibson assembly were from IDT. The sequences of all plasmids were confirmed by Sanger sequencing at the Genomics Shared Resource Facility (The Ohio State University). *E. coli* BL21 (DE3) cells were used for protein expression. Kanamycin (50 µg/ml) was added as needed.

Cells harboring ρ variants [all ρ variants are derivates of *E. coli* MG1655 ρ (NCBI ID: NP_418230.1) unless stated otherwise] were cultured in Terrific Broth (TB) at 37°C, and 0.5 mM isopropyl-1-thio-β-D-galactopyranoside (IPTG; Goldbio) was added when $OD_{600}$ reached ~0.5. After incubating for another 2.5 h at 37°C, the cells were collected by centrifugation at 8,000 x g for 8 min at 4°C.

For His-tagged ρ variants, the cell pellet was resuspended in Lysis buffer A (25 mM Tris-HCl, pH 7.5, 5 % (v/v) glycerol, 1 M NaCl, 0.5 mM tris(2-carboxyethyl)phosphine (TCEP), 1 x ProBlock Gold 2D Protease Inhibitor Cocktail (EDTA Free; Goldbio) and opened by sonication. The cell lysate was cleared by centrifugation (20,000 × g) for 30 min at 4°C. The supernatant was passed through a 0.45 µm filter and applied to HisTrap HP column (Cytiva). A linear gradient with Ni-A buffer (20 mM Tris-HCl, pH 7.5, 5% glycerol, 350 mM NaCl, 0.5 mM TCEP) and Ni-B buffer (Ni-A supplied with 1 M imidazole) were used for elution. The eluted protein was loaded onto HiTrap Heparin HP column (Cytiva) and eluted with a linear gradient of NaCl (0 - 1 M) in Hep-A buffer (20 mM Tris pH 7.5, 5% glycerol, 0.5 mM TCEP).

For tag-free ρ, cell pellet was resuspended in Lysis buffer B (50 mM Tris-HCl, pH 7.5, 10% glycerol, 100 mM KCl, 1 mM DTT, 1 x ProBlock Gold 2D Protease Inhibitor Cocktail (EDTA Free). After sonication, cell lysate was cleared by centrifugation and loaded onto HiTrap Heparin HP column. Protein was eluted as above and loaded onto Sephacryl S-300 HR (Cytiva), preequilibrated in 20 mM Tris pH 7.5, 5% glycerol, 0.5 mM TCEP, and 400 mM NaCl.

All ρ variants were dialyzed against dialysis buffer (10 mM Tris-HCl, pH 7.5, 5% glycerol, 500 mM NaCl, 0.5 mM TCEP) and flash frozen in liquid nitrogen.

The ρ proteins used for structural analysis by single partical cryoEM were purified as described above with the following variations. ρ proteins were expressed in *E. coli* RIL (BL21) cells (Novagen) and a final size-exclusion purification step was incorporated. Here, ρ mutants or ρ WT were loaded to a Superdex 200 column (Cytiva) equilibrated in SEC buffer (10 mM Tris pH 7.5, 200 mM NaCl, 1 mM DTT). Fractions corresponding to the peaks were combined and concentrated. ρ proteins were flash frozen in liquid nitrogen and stored at -80°C until further use.

Cells harboring *Pseudomonas aeruginosa* ρ (Pae-ρ) were cultured in TB at 37$^{o}$C, and 0.2 mM IPTG was added when $OD_{600}$ reached ~0.5 for overnight induction at 18$^{o}$C. The harvested cell pellet was resuspended in Lysis buffer C (40 mM Tris-HCl, pH 8, 5 % (v/v) glycerol, 0.5 M NaCl, 5 mM β-mercaptoethanol (β-ME), 1 x ProBlock Gold 2D Protease Inhibitor Cocktail (EDTA Free; Goldbio) and opened by sonication. The cell lysate was cleared by centrifugation (20,000 × g) for 30 min at 4$^{o}$C. The supernatant was passed through a 0.45 µm filter and applied to HisTrap HP column (Cytiva). A linear gradient with Ni-C buffer (40 mM Tris-HCl, pH 8, 5% glycerol, 300 mM NaCl, 5 mM β-ME, 0.1 mM PMSF) and Ni-D buffer (Ni-A supplied with 1 M imidazole) were used for elution. The eluted protein was loaded onto Resource Q column (Cytiva) after 1.5 times dilution with Q-A buffer (25 mM Tris pH 8, 5% glycerol, 5 mM β-ME). A linear gradient between Q-A and Q-B buffer (Q-A buffer plus 1 M NaCl) was applied. The fractions containing Pae-ρ were pooled and digested with ULP1 overnight at 4$^{o}$C. The next day, the sample was passed through HisPur Ni-NTA Resin (ThermoFisher) to remove the His-SUMO tag and ULP1. The untagged Pae-ρ was loaded onto HiTrap Heparin HP column and eluted with a linear gradient of NaCl (0 - 1 M) in Q-A buffer. The purity of Pae-ρ is assessed by mass spectrometry.

### Sample preparation for cryoEM

Before grid preparation, all ρ variants were applied to a Superdex 200 Increase 3.2/300 column (Cytiva) equilibrated with 10 mM Tris pH 7.5, 200 mM NaCl, 2 mM DTT and fractions of ρ were pooled and concentrated.

For ρ G150D and G152D, proteins (2.9 mg/ml) were incubated with 2.2 mM ADP-BeF for 5 min at room temperature. Subsequently, 3.8 µl of the ρ-ADP-BeF mix was applied to glow-discharged Quantifoil R1.2/1.3 holey carbon grids and plunged into liquid ethane using a Vitrobot Mark IV (Thermo Fisher) set at 10 °C and 100 % humidity. For $\rho^{pppGpp}$ and $\rho^{ADP}$, WT ρ (7.8 mg/ml) was incubated with 10 mM pppGpp or ADP, respectively, for 5 min at room temperature. 3.8 µl of the ρ-nucleotide mix was applied to glow-discharged Quantifoil R1.2/1.3 holey carbon grids and plunged into liquid ethane using a Vitrobot Mark IV (Thermo Fisher) set at 10 °C and 100 % humidity.

### CryoEM data acquisition and processing

CryoEM data were acquired on a FEI Titan Krios G3i TEM operated at 300 kV equipped with a Falcon 3EC direct electron detector. Movies were taken for 40.57 s accumulating a total electron dose of ~ 40 e-/Å2 in counting mode distributed over 33 fractions at a nominal magnification of 96,000x giving a calibrated pixel size of 0.832 Å/px.

All image analysis steps were done with cryoSPARC[1]. Movie alignment was done with patch motion correction generating half-binned micrographs, followed by CTF estimation with Patch CTF. Micrographs for ρ mutants G150D and G152D unexpectedly revealed filamentous organization of ρ besides hexamer particles. Class averages of manually selected particle images were used to generate an initial template for reference-based particle picking from 2,511 micrographs for the ρ-G150D sample. 495,704 particle images were extracted with a box size of 192 px and Fourier-cropped to 96 px for initial analysis. Reference-free 2D classification was used to select 168,536 particle images of filamentous ρ for further analysis. Helix refinement applying a helical rise of 200 Å was used to generate an initial 3D reconstruction of the filament from a small subset of the data set. To obtain a better resolved reconstruction, all selected particle images were refined against this reference low-pass filtered to 30 Å applying the non-uniform (NU) refinement routine. Local processing was allowed to start at a resolution of 8 Å. Particle images were re-extracted with a box size of 384 px after local motion correction and subjected to helix refinement giving a reconstruction at ~ 3.5 Å resolution. Another iteration of reference-free 2D classification was applied to select the best class averages, followed by local CTF refinement and helix refinement to generate the final reconstruction at ~ 3.3 Å resolution. Data for the ρ-G152D sample was analysed accordingly, giving a final reconstruction at ~ 3.3 Å resolution.

Since WT ρ samples did not reveal any filamentous particles, ab initio reconstruction was used instead of helix refinement to generate an initial reference for 3D refinement by NU refinement. To account for sub-stoichiometric composition of ρ oligomers and isolate hexamers, 3D variability analysis (3DVA) was conducted. Final data sets of 200,796 and 314,323 particle images were selected for NU refinement, giving reconstructions of wildtype ρ at 3.0 Å and 2.6 Å in presence of pppGpp and ADP, respectively.

### Model building, refinement, and analysis

Coordinates of ρ (PDB ID: 6WA8 [open ring][2]) were docked into the cryoEM maps using Coot (version 0.8.9)[3]. ρ subunits were manually adjusted to fit the cryoEM density. Manual model building alternated with real space refinement in PHENIX (version 1.20.1)[4]. Data collection and refinement statistics are provided in Table S1. Structure figures were prepared with ChimeraX (version 1.6.1)[5,6].

#### *In vitro* pelleting assay

1 µM purified ρ in PN23 buffer (10 mM Tris-HCl, pH7.4, 0.5 mM TCEP, 12 mM $MgCl_2$, 230 mM NaCl) was cleared at 10,000 x g for 5 min at 20°C immediately before pelleting assay. Nucleotides were mixed with the cleared ρ and incubated at room temperature for 10 min. Then 40 µL of reaction was centrifuged at 55,000 rpm (~100,000 x g) in a TLA100 rotor (Beckman Coulter) for 20 min at 20°C. The supernatant was removed and pellet was soaked in 40 µL of RRB buffer (10 mM Tris-HCl, pH7.4, 1 mM DTT, 500 mM NaCl, 2 M Urea) for 1 h at room temperature before resuspending. The samples were analyzed on SurePAGE 4-12% gel (GenScript) and stained with GelCode Blue Safe Protein Stain (Thermo).

#### *In vivo* pelleting assay

To determine the influence of translation-inhibiting antibiotics on ρ filament formation, *E. coli* Δ*rfaH rho*$^+$ strain (IA228; all strains used in this study are listed in Table S2) was cultured in LB at 37°C to $OD_{600}$ ~0.5. Then antibiotics were added at the following final concentrations: 30 µg/mL nalidixic acid, 100 µg/mL mupirocin, 50 µg/mL retapamulin, or 100 µg/mL rifampicin. Ethanol was added to 0.4% (v/v) as a negative control. After 30 min incubation at 37°C, the cells were collected by centrifugation. To further test the influence of stress removal on ρ filament, after 30 min exposure to mupirocin, the cells were washed three times, reinoculated into the same volume of fresh LB, and incubated at 37°C. The cells were collected after 40 min recovery. The growth curves for mupirocin recovery were recorded with EPOCH 2 microplate reader (BioTek) (Extended Data Fig. 7).

To test the effects of cellular starvation of $Mg^{2+}$ on formation of ρ polymers, we adapted a previously published protocol[7]. A modified MOPS medium (MM medium) was used: MOPS medium lacking $Mg^{2+}$ and $Ca^{2+}$ ions[8] was supplied with 0.1% casamino acids and 0.2% glucose. Strain IA228 was cultured overnight in MM medium supplied with 10 mM $MgCl_2$. The next day, cells were reinoculated into fresh MM medium plus 10 mM $MgCl_2$, and allowed to grow to $OD_{600}$ ~0.2 at 37°C. Then cells were washed 3 times with MM medium to remove $Mg^{2+}$. Cells were resuspended in the same volume of fresh MM medium supplied with either 0 mM or 10 mM $MgCl_2$. Cells were pelleted after 2 and 3 h incubation.

Collected cells were opened using sonication in PN23 buffer supplied with 1 x ProBlock Gold 2D Protease Inhibitor Cocktail (EDTA Free; Goldbio). Cell lysate was centrifuged at 1,000 x g for 10 min at 4°C, and the supernatant was passed through a 0.45 μm filter. The cell-free cell lysate was centrifuged at 55,000 rpm in TLA100 rotor for 30 min at 4$^{o}$C. The supernatant was removed, and the pellet was soaked in 30 μL of RRB buffer for 1 h before resuspending. The samples were analyzed by Western blotting with anti-ρ polyclonal antibodies (a gift from Evgeny Nudler, New York University).

### Western blot analysis

Following separation in SurePAGE gels, the proteins were transferred to nitrocellulose membrane (Bio-Rad) by electrophoresis with 100 V for 1 h on ice in Transfer buffer (25 mM Tris, pH 8.3, 192 mM glycine, and 20% methanol). After the transfer step, the membrane was blocked in Blocking buffer (25 mM Tris-HCl, pH 7.4, 150 mM NaCl, 0.1% Tween-20, and 5% (w/v) Blotting Grade Blocker Non-Fat Dry Milk (Bio-Rad)) for 1 h at room temperature. The membrane was washed twice with TBST (25 mM Tris-HCl, pH 7.4, 150 mM NaCl, 0.1% Tween-20) followed by incubation with anti-ρ polyclonal antibodies (1:10,000) in Blocking buffer overnight at 4$^{o}$C. The next day, the membrane was washed five times with TBST and incubated with secondary antibody (1:10,000; Goat Anti-Rabbit IgG (H + L)-HRP Conjugate (Bio-Rad)) in Blocking buffer for 1 h at room temperature. The membrane was then washed five times with TBST. Finally, the membrane was incubated with Clarity Max Western ECL Substrate (Bio-Rad) and imaged with ChemiDoc XRS+ System. Quantification was done with Image Lab v6.1 (Bio-Rad). The means and p-value were calculated in Excel (Microsoft).

### *Ex vivo* crosslinking

To visualize the filament formation *in vivo*, *E. coli* BL21 (DE3) cells harboring plasmids encoding $His_8$-tagged ρ variants were cultured in 50 mL of LB at 37$^{o}$C to $OD_{600}$~0.7 and protein expression was induced with 1 mM IPTG for a 2 h induction at 37$^{o}$C. Cells were pelleted by centrifugation and washed twice with 1 x PBS (10 mM phosphate buffer, 2.7 mM KCl and 137 mM NaCl, pH 7.4). After resuspending in 1 x PBS, 1 mM bismaleimidoethane (BMOE; Thermo) was added. Crosslinking was done for 20 min at room temperature in the dark, followed by quenching with 280 mM β-ME for 10 min. The cells were collected and resuspended in sonication buffer (1 x PBS, 400 mM NaCl, 2 mM DTT, 0.1 mM PMSF, and 2 M Urea). After sonication, the cell lysate was cleared at 20,000 x g for 10 min. Proteins were resolved on a SurePAGE 4-12% gel, stained with NTA-Atto 550 (Sigma), and visualized on Typhoon 5 (Cytiva) with Cy3 mode.

### *In vitro* crosslinking

Nucleotides were added at concentrations indicated in figure legends to 1 µM purified $\rho^{X1}$ in PN23 buffer and incubated for 10 min at room temperature. Then 0.5 mM BMOE was added, followed by 10 min incubation at room temperature in the dark. The reaction was quenched with 280 mM β-ME for 5 min. To assay for filament dispersal, filaments were first formed with ADP/ppGpp in PN23 supplied with an additional 10 mM $MgCl_2$. Then, ATP-γS was added, followed by 30 min incubation at room temperature. Then samples were crosslinked with BMOE and quenched as above.

After separation in SurePAGE 4-12% gels, His-tagged ρ was stained with NTA-ATTO 550, and tag-free ρ was stained with GelCode Blue Safe Protein Stain. Gels were scanned and quantified using Typhoon 5 and ImageQuant v5.2. The means and standard deviation (SD) were calculated in Excel (Microsoft).

### Sucrose gradient

Sucrose gradients were prepared as described in[9]. Briefly, different sucrose density layers were prepared in 5% sucrose steps, filtered through a 0.45 µm filter, and sequentially layered into 13.5 mL open-top thin-wall polypropylene tubes (Beckman Coulter), starting with the highest density; each layer was incubated at -80°C for 15-30 min before addition of the next.

For testing cell extracts, *E. coli* Δ*rfaH* strains with *rho*$^+$ (IA228) and *rhoG150D* (IA305) chromosomal alleles were cultured in LB at 37°C for 24 h. Cells were collected by centrifugation and opened using sonication in PN23 buffer supplied with 1 x Protease Inhibitor Cocktail. Cell lysate was centrifuged at 1,000 x g for 10 min at 4°C, and the supernatant was passed through a 0.45 µm filter. Then 100 µL of the cell-free lysate was loaded onto sucrose gradient.

To analyze purified ρ, 1 µM ρ was incubated with indicated nucleotides for 10 min at room temperature. Then 100 µL of the reaction was loaded onto sucrose gradient.

Ultracentrifugation of sucrose gradients was performed at 110,000 x g in SW41Ti rotor (Beckman Coulter) for 16 h at 4°C. Fractioning of the gradient was done by taking aliquots from the top of the gradient, and the pellet was resuspended in RRB buffer. The samples were analyzed by Western blotting.

### Tryptophan fluorescence

Tryptophan fluorescence spectroscopy was performed using F-7000 Fluorescence Spectrophotometer (Hitachi) at room temperature. The excitation wavelength was set at 280 nm and the emission spectra were recorded from 300 to 390 nm with a 5 nm slit width of excitation and emission. The scan speed was 240 nm/min. The samples were prepared in PN23 buffer. Prior to recording the spectra, 1 µM ρ was incubated with 0.25 mM nucleotides at room temperature for 3 min.

#### *In vivo* ppGpp sensitivity assays

Three assays were performed to compare the effect of (p)ppGpp on the WT and G150D *rho* strains. First, the *relA* deletion strains carrying either WT (IA793) or G150D (IA791) *rho* variants were transformed with plasmids expressing either WT or catalytically-dead D275G (p)ppGpp synthetase RelA from an IPTG-inducible promoter[10], plated on LB/Carb in the absence of inducer, and incubated at 37°C for 16 hours (Fig. 4b). Second, the effect of *relA* deletion on growth of WT and G150D strains was compared in liquid growth assays at 37°C (Extended Data Fig. 5d). Third, strains with WT (IA228), G150D (IA305), and a *rho*-down [an IS2 insertion in *rhoL* (IA306)] alleles were streaked out on LB plates with or without 25 mg/L mupirocin and incubated overnight at 37°C (Extended Data Fig. 5e).

#### LigPlot$^+$

LigPLot$^+$ (v.2.2.8) is the software tool to run the LigPlot and DimPlot programs[11]. The LigPlot and DimPlot programs identify various types of interactions and generate schematic diagrams that show the contacts between protein-ligand and protein-protein residues, respectively. The diagrams visualizes salt bridges (red dashes), hydrogen bonds (green dashes) and hydrophobic contacts (red rays). For the characterization of inter-subunit contacts (Fig. 3e), hydrophobic contacts have been omitted for clarity.

#### Determinig hexamer geometry

For the calculation of hexamer geometry, as depicted in Fig. 1d, the "draw_rotation axis" script in PyMOL was utilized. The script calculates three values between two adjacent subunits: the rotation angle (β) with respect to the rotation axis, the length of the translation vector along the rotation axis (r, between A/B, B/C, C/D, D/E, and E/F), and p (between A/F). Additionally, it computes the distance between two adjacent subunits (designated as "g" for the A-F distance). Subsequently, these values were employed to calculate the upward rotation (α) between subunits. The presented values for α and r represent the means determined between A/B, B/C, C/D, D/E, and E/F.

#### ρ distribution analysis

The distribution of ρ was estimated using the ρ TIGR model (TIGR00767) in Annotree[12] with an e-value

$10^{-5}$ (Dataset 1a). The result was visualized in R (v. 4.2.2) with the Ggtree package[13]. Since phyla with many sequenced genomes are over-represented in the Annotree dataset, this dataset is not suitable for conservation analysis.

### ρ conservation

To compile a representative database for ρ conservation study. The GTDB[14] bacterial taxonomy list (Released April 08, 2022) was downloaded. Genomes labeled as "gtdb_type_species_of_genus" and "gtdb_representative" were kept. Genome assembly level was inspected. Only complete genomes were used to build the database. The new dataset covers 41 Phyla including 649 Genera. The protein sequences of all genomes were downloaded from NCBI. To identify ρ, Pfam model Rho_RNA_bind (PF07497) was searched against the representative database using hmmsearch (v. 3.3)[15] with a bit score of 27. The identified ρ-like proteins were further confirmed by searching NCBI HMM model of ρ (NF006886.1) against them using hmmsearch with cutoff 315. Finally, 632 ρ sequences are collected (Dataset 1b).

The length of ρ sequences ranges from 355 to 857 residues. Many of them have long insertions. Multiple sequence alignment (MSA) was done by Dialign v2.2.1[16] with default setting, which is suitable for detecting local homologies in sequences with low overall similarity. For comparision purpose, the MSA was indexed according to *E. coli* MG1655 ρ (NP_418230.1). The MSA is inspected manually in Jalview v2.11.2.7[17]. Sequence logo was generated using WebLogo (v. 3.7.8)[18].

### ρ clustering analysis

The flagellar export ATPase FliI was used as a decoy during clustering; FliI (NP_416451.1) has the highest bit score when searching ρ (NP_418230.1) against *E. coli* MG1655 genome using blastp. To collect decoy sequences, NP_416451.1 was used as a query to do blastp searching against the representative database with e-value $10^{-5}$. Two decoy sequences were randomly selected from each Phylum. A total of 42 decoy sequences were selected (Dataset 1c). An all-vs-all blastp was performed using BLAST+ (v. 2.9.0)[19] with an e-value of $10^{-10}$. A ρ similarity network was built based on e-value $10^{-99}$, which can separate decoy and ρ very well. Markov clustering was done based on the bit scores of all-vs-all blastp result in Cytoscape (v. 3.9.1)[20] with clusterMaker2 (v. 2.3.2)[21]. Among 41 ρ clusters, those with 10+ sequences were used for further analysis (Dataset 1d). The presence of IDR (intrinsically disordered regions) and prion-like domains were detected by metapredict v2.6[22] and PrionW[23], respectively.

### Table S1. CryoEM data collection, refinement, and validation statistics.

| Data collection and processing | | | | |
|---|---|---|---|---|
| | ρ$^{G150D}$ | ρ$^{G152D}$ | ρ$^{pppGpp}$ | ρ$^{ADP}$ |
| Microscope | FEI Titan Krios G3i | | | |
| Voltage [keV] | 300 | | | |
| Camera | Falcon 3EC | | | |
| Magnification (nominal) | 96,000x | | | |
| Pixel size at detector [Å/pixel] | 0.832 | | | |
| Total electron exposure [$e^-$/Å$^2$] | 42 | | | |
| Exposure rate [$e^-$/pixel/s] | 0.7 | | | |
| Frames collected during exposure | 33 | | | |
| Defocus range [µm] | 0.8 - 2 | | | |
| Automation software | EPU version 2.10 | | | |
| Micrographs | | | | |
| Collected | 2511 | 1642 | 1477 | 1599 |
| Used | 2511 | 1578 | 1467 | 1544 |
| Particle images | | | | |
| Total extracted | 495,704 | 428,112 | 641,595 | 602,082 |
| Final | 140,359 | 222,106 | 200,796 | 314,323 |
| Point-group or helical symmetry parameters | C1 | C1 | C1 | C1 |
| Resolution [Å] | | | | |
| Global | | | | |
| $FSC_{0.143}$[(a)] (unmasked/masked) | 3.9 / 3.3 | 3.8 / 3.3 | 3.5 / 3.0 | 3.1 / 2.6 |
| Local resolution range [Å$^2$] | 1.8 - 39 | 1.8 - 30 | 2.2 - 35 | 1.8 - 35 |
| Map sharpening *B* factor [Å$^2$] | -87.6 | -92.7 | -101.7 | -97.6 |
| Map sharpening methods | local B-factor | | | |
| Refinement software | | | | |
| Package | PHENIX version 1.20_4459 | | | |
| Routine | real.space.refine | | | |
| **Model composition** | | | | |
| | ρ$^{G150D}$ | ρ$^{G152D}$ | ρ$^{pppGpp}$ | ρ$^{ADP}$ |
| Model composition | | | | |
| Non-H atoms | 59.922 | 59.922 | 19.9007 | 19.912 |
| Protein residues | 7.542 | 7.542 | 2.508 | 2.508 |
| RNA residues | - | - | - | - |
| $Mg^{2+}$ ions | 18 | 18 | -1 | 4 |
| ADP | 18 | 18 | - | 6 |
| BeF | - | - | - | - |
| pppGpp | - | - | 4 | - |
| **Model Refinement** | | | | |
| Model-Map scores | | | | |
| CC[(b)] (mask) | 0.86 | 0.87 | 0.82 | 0.85 |
| CC (volume) | 0.86 | 0.87 | 0.82 | 0.85 |
| Average grouped B factors [Å$^2$] | | | | |
| Overall | 133 | 129 | 106 | 117 |
| Protein | 133 | 129 | 106 | 117 |
| $Mg^{2+}$ ions | 123 | 138 | 37 | 70 |
| ADP | 144 | 137 | - | 134 |
| pppGpp | - | - | 99 | - |
| Rmsd[(c)] from ideal values | | | | |
| Bond lengths [Å] | 0.004 | 0.003 | 0.005 | 0.004 |
| Bond angles [°] | 0.819 | 0.807 | 0.625 | 0.688 |
| **Validation[(d)]** | | | | |
| MolProbity score | 2.05 | 2.02 | 1.62 | 1.52 |
| CaBLAM outliers [%] | 0.96 | 0.72 | 0.89 | 0.97 |
| Clashscore | 13.09 | 15.02 | 12.97 | 9.72 |
| Poor rotamers [%] | 2.78 | 2.78 | 0.33 | 1.02 |

| | | | | |
|---|---|---|---|---|
| Cβ deviations | 0.0 | 0.0 | 0.0 | 0.0 |
| EMRinger score | 1.79 | 1.70 | 1.71 | 2.21 |
| Ramachandran plot | | | | |
| Favored [%] | 97.60 | 98.56 | 98.12 | 98.56 |
| Allowed [%] | 2.40 | 1.44 | 1.88 | 1.44 |
| Outliers [%] | 0.0 | 0.0 | 0.00 | 0.0 |
| Ramachandran *Z*-score (rmsd) | | | | |
| Overall | 0.90 (0.10) | 1.21 (0.10) | 0.89 (0.17) | 0.85 (0.17) |
| Helices | 1.27 (0.09) | 1.33 (0.09) | 1.31 (0.16) | 1.14 (0.16) |
| Sheets | 0.28 (0.15) | 1.11 (0.15) | 0.12 (0.26) | 0.86 (0.28) |
| Loops | 0.17 (0.11) | 0.14 (0.11) | 0.03 (0.20) | -0.11 (0.19) |
| Data deposition | | | | |
| Reconstruction (EMDB) | EMD-18132 | EMD-18133 | EMD-18131 | EMD-18130 |
| Coordinates (PDB) | 8Q3P | 8Q3Q | 8Q3O | 8Q3N |

[a] FSC, Fourier shell correlation
[b] CC, correlation coefficient
[c] Rmsd, root-mean-square deviation
[d] Using MolProbity[24].

### Table S2. Plasmids and strains used in this study.

| Plasmids | Key features | Source |
|---|---|---|
| pET24-ρ | Tagless wild type ρ under T7 promoter | [25] |
| pIA1301 | T7 promoter- ρ-$His_6$ | This study |
| pIA1307 | T7 promoter- ρ [G150D]-$His_6$ | This study |
| pIA1510 | T7 promoter- ρ [S84C]-$His_8$ | This study |
| pIA1511 | T7 promoter- ρ [G150D S84C]-$His_8$ | This study |
| pIA1513 | T7 promoter- ρ [E106C E375C]-$His_8$ | This study |
| pIA1514 | T7 promoter- ρ [S84C M405C]-$His_8$ | This study |
| pIA1515 | T7 promoter- ρ [G150D S84C M405C]-$His_8$ | This study |
| pIA1516 | T7 promoter- ρ [G150D E106C E375C]-$His_8$ | This study |
| pIA1539 | T7 promoter- ρ [M405C] | This study |
| pIA1540 | T7 promoter- ρ [S84C M405C] | This study |
| pIA1634 | T7 promoter-$His_{10}$-SUMO-*P. aeruginosa* ρ | This study |
| pR1-1His | Wild type RelA under IPTG-inducible promoter | [10] |
| pR1-1His(D275G) | Inactive RelA mutant under IPTG-inducible promoter | [10] |
| **Strains** | **Genotype** | **Source** |
| IA228 | MG1655 Δ*rfaH* | [26] |
| IA305 | MG1655 Δ*rfaH* Δ*rac rho*G150D | [26] |
| IA306 | MG1655 Δ*rfaH* Δ*rac rhoL*-ΩIS2-*rho* | [26] |
| IA539 | MG1655 Δ*rfaH* Δ*rac* | This study |
| IA791 | MG1655 Δ*rfaH* Δ*rac rho*G150D *relA*::Kn | This study |
| IA793 | MG1655 Δ*rfaH* Δ*rac relA*::Kn | This study |

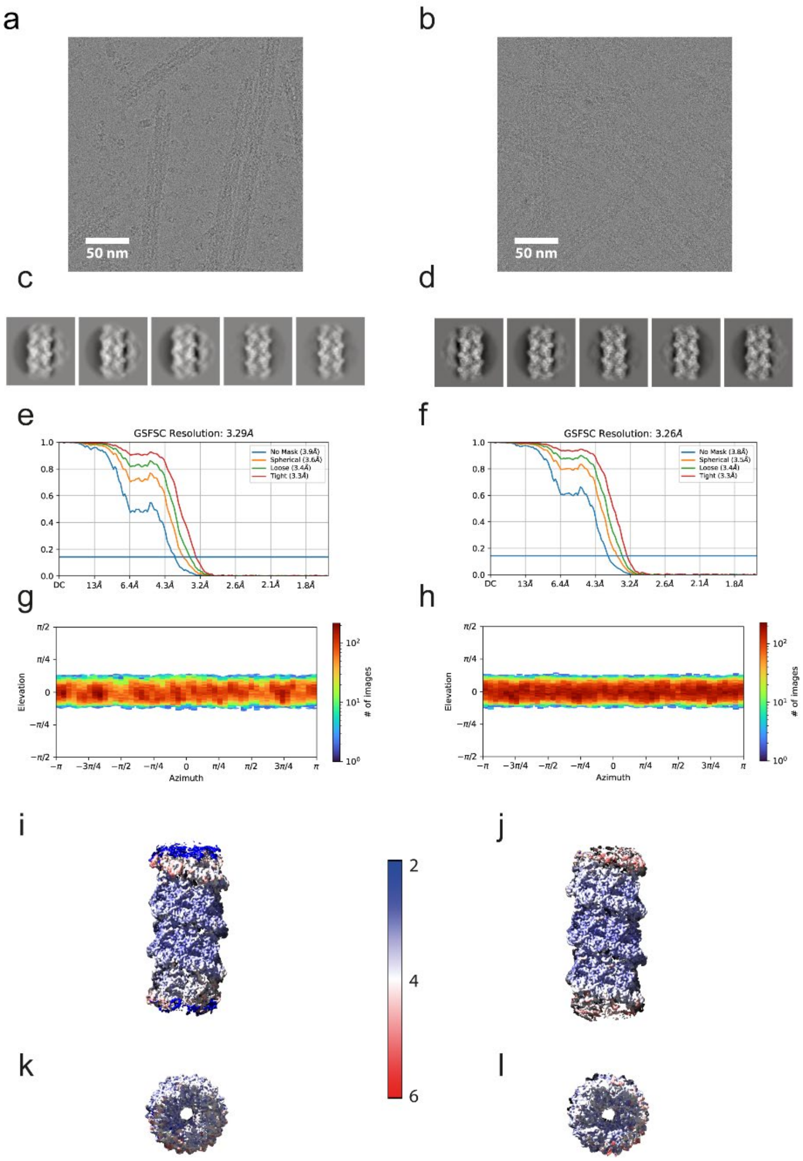


**Extended Data Fig. 1. CryoEM analysis of ρ mutants G150D (left) and G152D (right). (a/b)** representative cryoEM micrographs, scale bar corresponds to 50 nm. **(c/d)** selected 2D class averages after reference-free 2D classification. **(e/f)** Gold-standard fourier-shell-correlation curves and **(g/h)** viewing direction distribution after helix refinement. 3D reconstruction colored by local resolution viewed along the helical axis **(i/j)** and rotated by 90 degrees **(k/l)**.

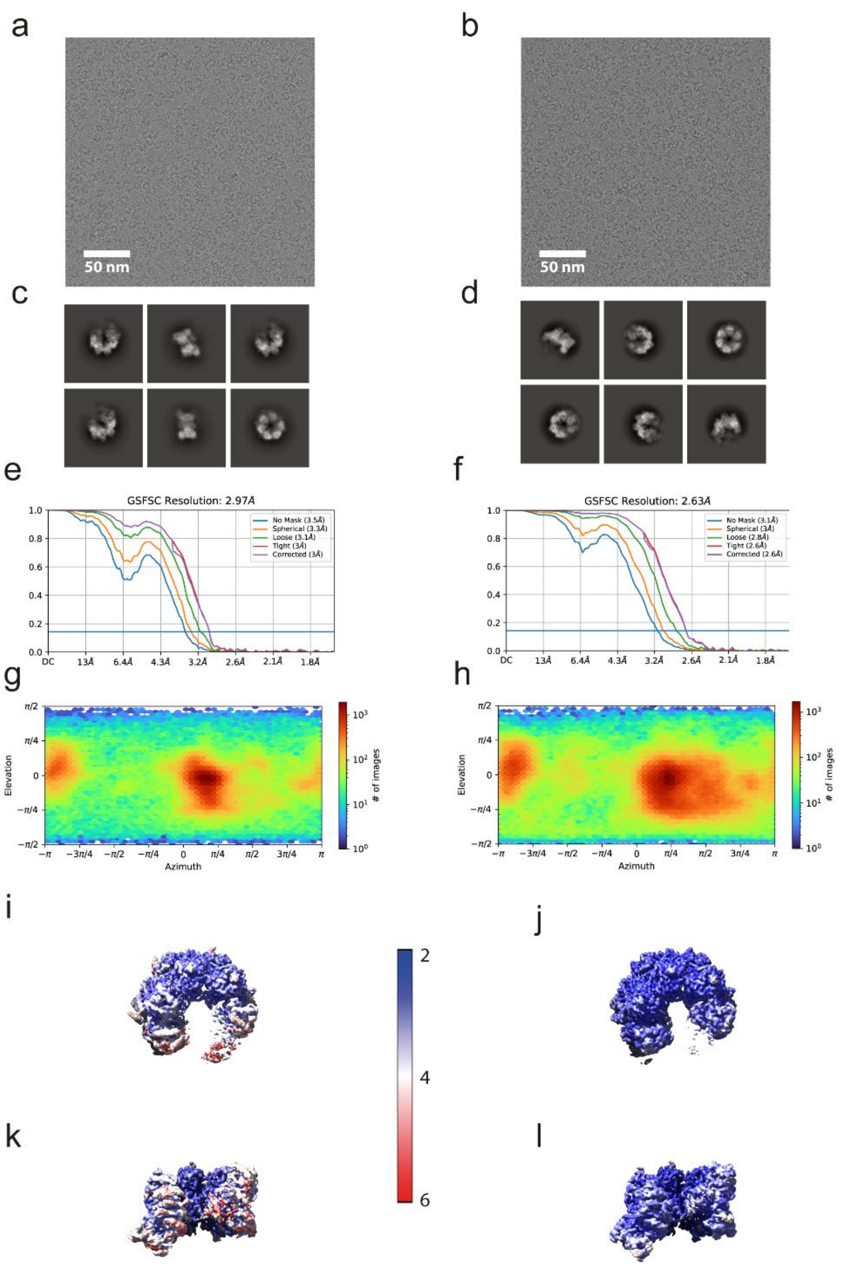


**Extended Data Fig. 2. CryoEM analysis of WT ρ•pppGpp (left) and ρ•ADP (right). (a/b)** representative cryoEM micrographs, scale bar corresponds to 50 nm. **(c/d)** selected 2D class averages after reference-free 2D classification. **(e/f)** Gold-standard fourier-shell-correlation curves and **(g/h)** viewing direction distribution after non-uniform refinement. 3D reconstruction colored by local resolution from top view **(i/j)** and side view (rotated by 90 degrees) **(k/l)**.

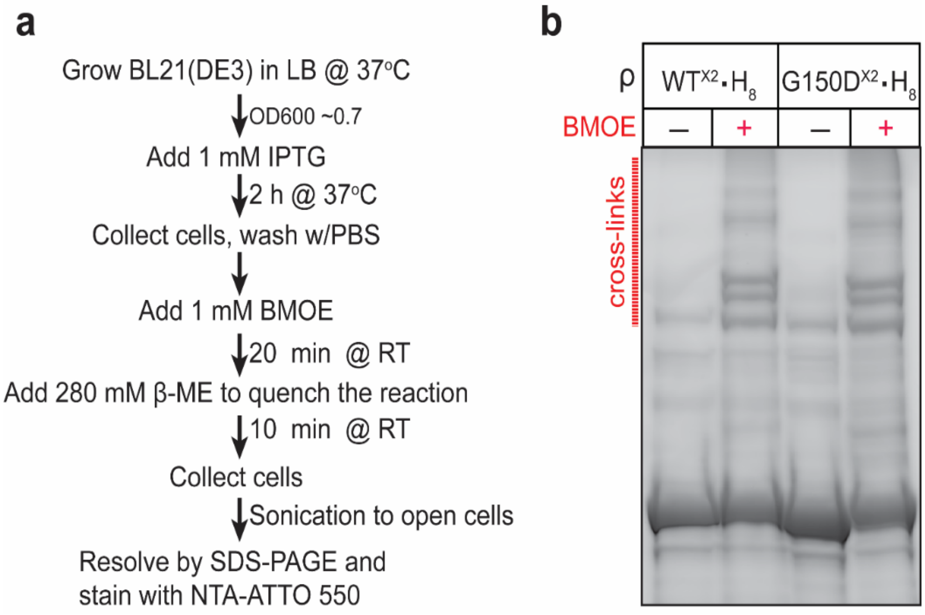


**Extended Data Fig. 3**. **BMOE mediated *ex vivo* cysteine cross-linking with the C106/C375 sensor pair (X2).** **a,** Schematic of the experiment with C-terminally $His_8$-tagged ρ, WT or G150D, bearing X2 substitutions. RT, room temperature. **b**, Crosslinking products were detected by LDS-PAGE and in-gel fluorescence using His-tag specific NTA-ATTO 550 stain.

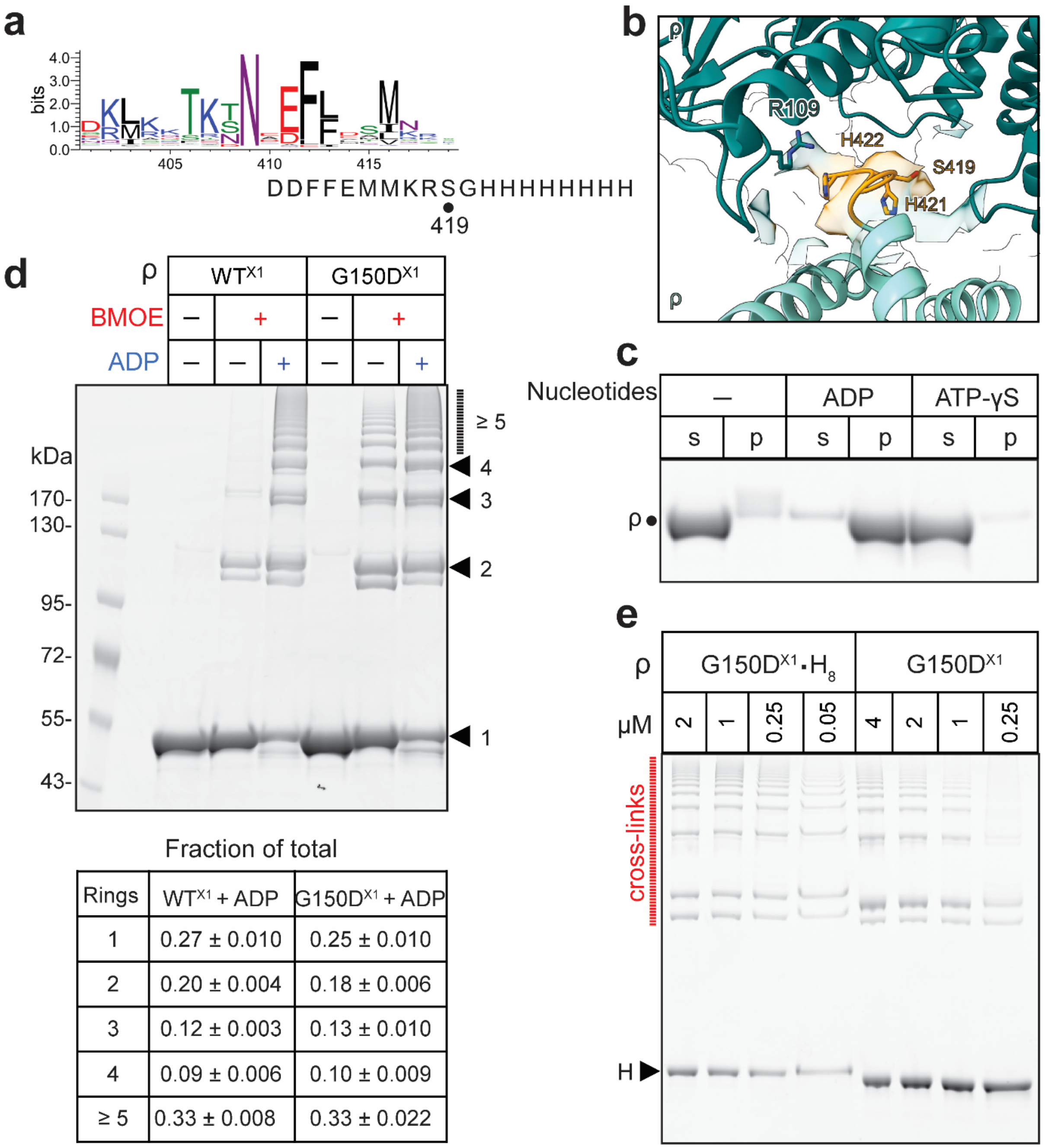


| Rings | WT^X1 + ADP | G150D^X1 + ADP |
|---|---|---|
| 1 | 0.27 ± 0.010 | 0.25 ± 0.010 |
| 2 | 0.20 ± 0.004 | 0.18 ± 0.006 |
| 3 | 0.12 ± 0.003 | 0.13 ± 0.010 |
| 4 | 0.09 ± 0.006 | 0.10 ± 0.009 |
| ≥ 5 | 0.33 ± 0.008 | 0.33 ± 0.022 |

**Extended Data Fig. 4. His tag is not essential for filamentation but promotes oligomerization at lower ρ concentrations. a,** The conservation of the ρ C-terminus across the bacterial kingdom. The His-tagged ρ used in this study has 8 histidines after residue S419. **b**, The view of His tag position in the structure, which is highlighted in orange. **c,** 1 μM of tag-free WT ρ is pelleted in the presence of 2.5 mM ADP, but not 2.5 mM ATP-γS. s, supernatant; p, pellet. **d**, ADP-induced filamentation of tag-free WT and G150D ρ variants bearing X1 Cys substitutions. 1.5 μM ρ and 2.5 mM ADP were used in this BMOE crosslinking assay. Abundance of hexamers (1) and higher-order oligomers was quantified in three independent experiments; the results are shown as mean ± SD. **e**, Filamentation of G150D•$H_8$ is observed at 50 nM, whereas tag-free G150D forms filaments only above 1 μM.

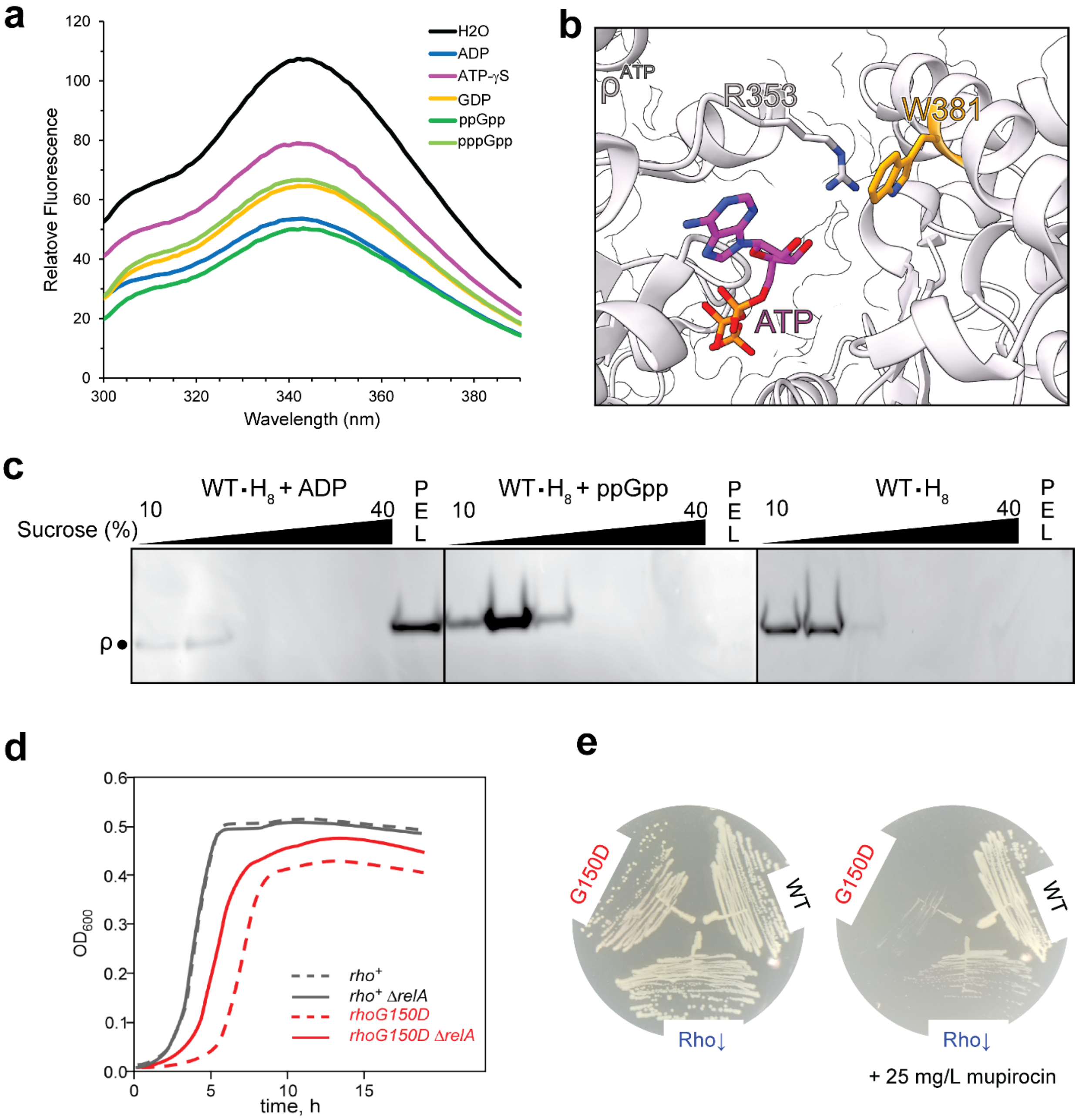


**Extended Data Fig. 5**. **ppGpp binds to ρ to promote oligomerization *in vitro* and compromises the survival of G150D *rho* strain. a,** Tryptophan fluorescence spectra show the quenching of W381 upon nucleotide binding**. b,** W381 is located near the nucleotide-binding pocket. **c,** Separation of ADP/ppGpp-induced higher-order oligomers on sucrose gradients. **d,** Growth of WT and G150D *rho* strains carrying *relA*+ or their Δ*relA* derivatives; the Δ*relA*::Kan was moved by P1 transduction from the Keio collection strain[27]. Overnight cultures were diluted 1:100 into fresh LB and grown at 37°C. The growth curves were recorded with EPOCH 2 microplate reader (BioTek). **e**, Growth of *rhoG150D* is severely inhibited by mupirocin (MUP), which induces stringent response and (p)ppGpp accumulation[28]. The inhibitory effect of

MUP is less pronounced in a strain in which ρ levels are reduced to 40% by an IS2 insertion into the *rho* leader region[26]. All experiments were done in triplicates.

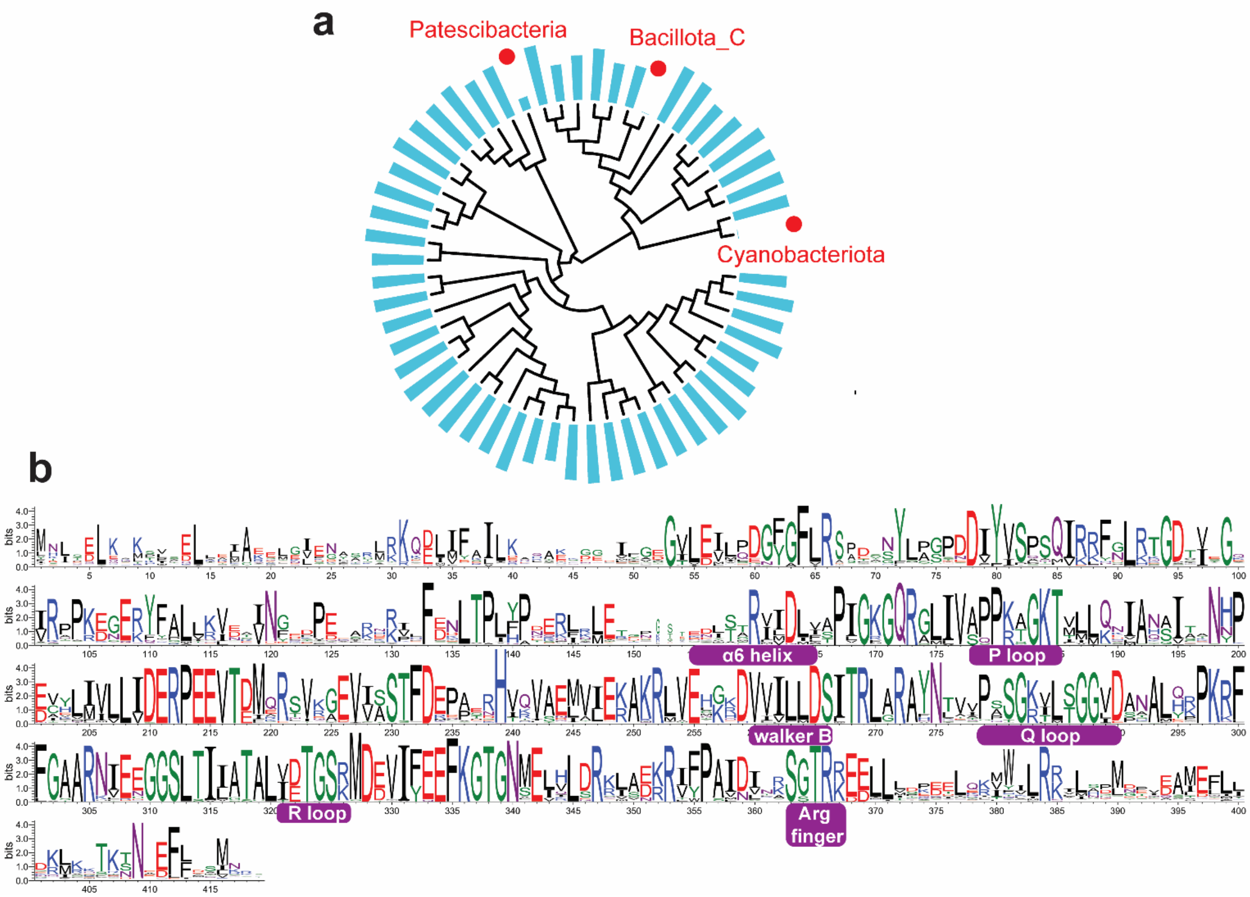


**Extended Data Fig. 6**. **Conservation of ρ**. **a,** ρ distribution on Phylum-level resolution. Bars represent the percentage of genome hits in each Phylum calculated from the Annotree dataset. Phyla with < 10 genomes are removed (see Dataset 1 and Methods for details). Three Phyla indicated in red show low or no ρ hits. **b**, ρ sequence conservation in bacteria. Sequence logos were generated by WebLogo (version 3.7.8).

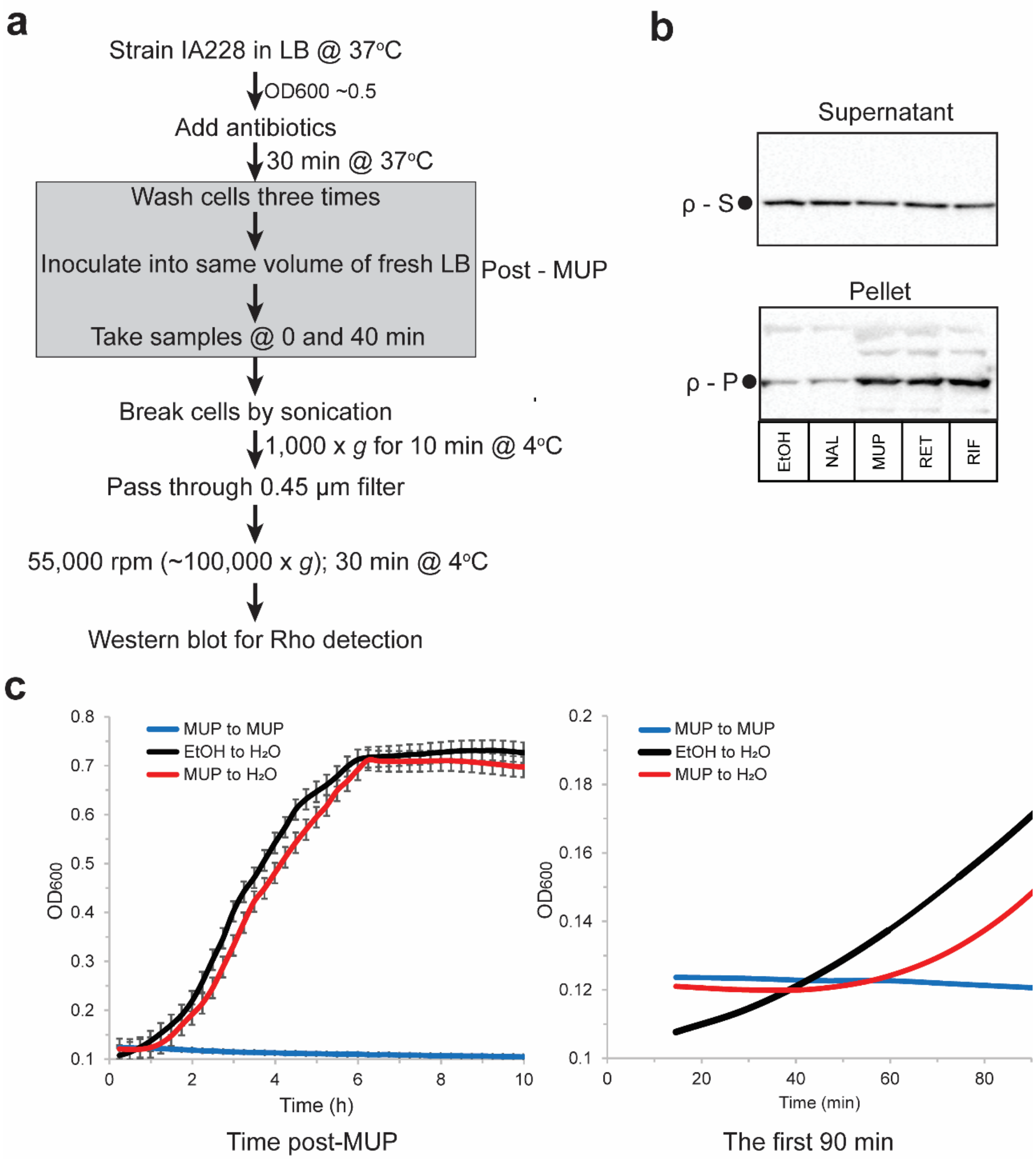


**Extended Data Fig. 7**. **Antibiotic stresses affect ρ filamentation *in vivo***. **a,** Schematic illustration of *in vivo* pelleting assay. **b**, Representative western blot result. After ultracentrifugation, ρ in the pellet (ρ-P) and supernatant (ρ-S; samples of the supernatant were diluted 15 times before loading) were resolved and detected by Western blot. The fraction of ρ in the pellet was calculated as 100 × [ρ-P/(ρ-P + ρ-S)]. EtOH, the same amount of ethanol was served as control; NAL, nalidixic acid; MUP, mupirocin; RET, retapamulin; RIF, Rifampicin. **c,** Growth curves of the MUP recovery. MUP/EtOH treated cells were washed and reinoculated into fresh medium supplied with MUP or the same amount of autoclaved water. Right, overall view of the growth curves. Cells treated with MUP were resuspended in fresh medium supplied with MUP (blue line, MUP to MUP). Cells treated with ethanol were resuspended in medium supplied with the same amount of water (black line, EtOH to $H_2O$). The red line (MUP to $H_2O$) represents the MUP treated cells recovering in fresh medium. Left, a zoom-in view to show the recovery stage. The error bars are omitted for clarity. Experiments were done in triplicates.

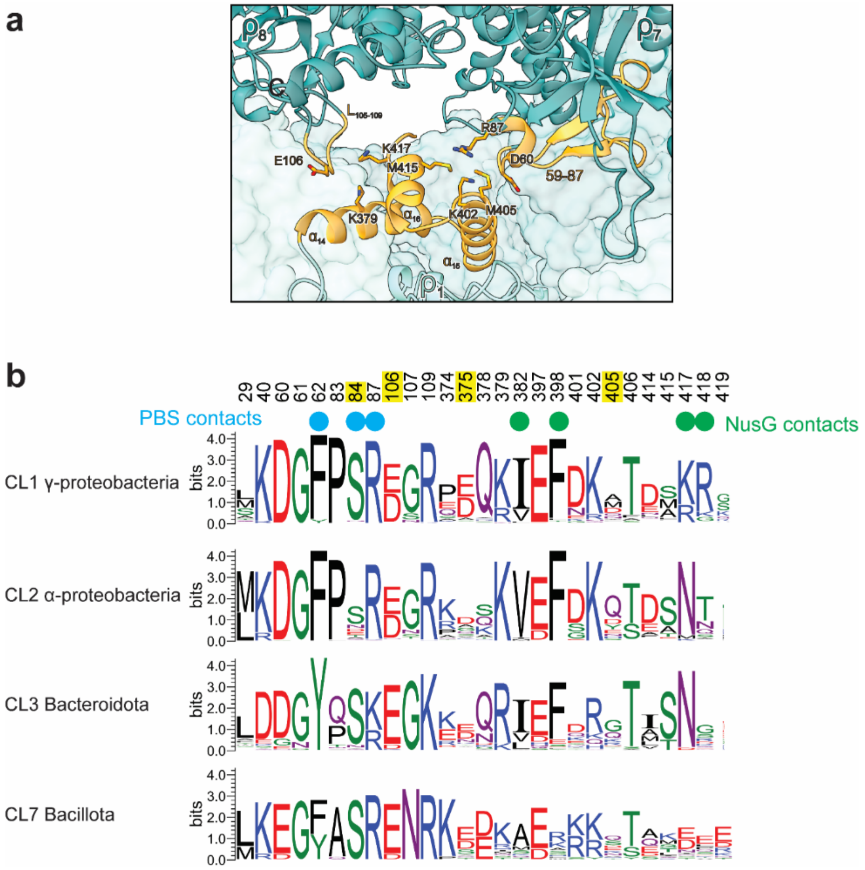


**Extended Data Fig. 8**. **Conservation of the filament interface**. **a,** Structural view of the filament interface. Interaction residues are denoted in orange. **b,** Sequence logos showing the conservation of filament interface in four of the ρ clusters. The four residues mutated to Cys are highlighted in yellow. PBS (cyan dots) and NusG (green dots) contact residues overlapped with the filament interface.

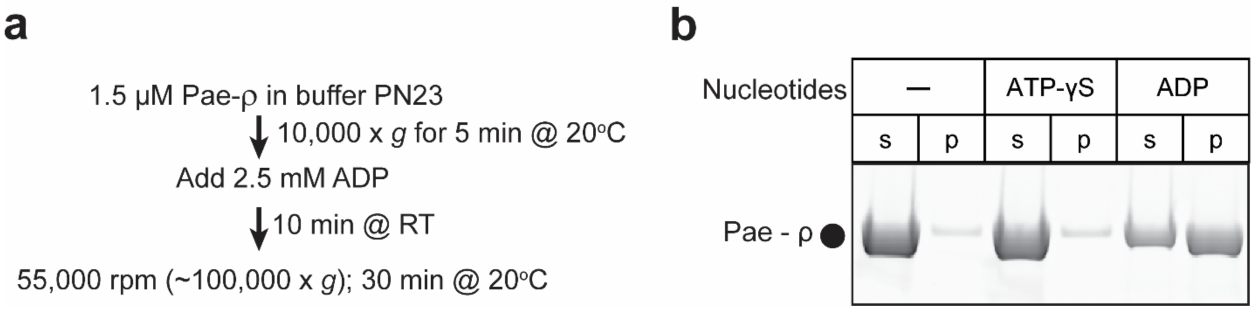


**Extended Data Fig. 9. *P. aeruginosa* ρ aggregates in the presence of ADP**. Pae-ρ and Eco-ρ belong to CL1 (Fig. 6) and neither protein has an IDR or prion-like domain. Pelleting assays show that Pae-ρ can polymerize upon incubation with ADP, but not ATP-γS; the same behavior was observed with Eco-ρ (Fig. 2a).
